# A Novel Index for Predicting Health Status Using Species-level Gut Microbiome Profiling

**DOI:** 10.1101/2020.02.24.962100

**Authors:** Vinod K. Gupta, Minsuk Kim, Utpal Bakshi, Kevin Y. Cunningham, John M. Davis, Konstantinos N. Lazaridis, Heidi Nelson, Nicholas Chia, Jaeyun Sung

## Abstract

The development of a biologically-interpretable and robust metric that provides clear insight into the general health status (i.e. healthy or non-healthy) of one’s gut microbiome remains an important target in human microbiome research. We introduce the Gut Microbiome Health Index (GMHI), a mathematical formula that determines the degree to which a gut microbiome profile reflects good or adverse health. GMHI was formulated based on microbial species specific to healthy gut ecosystems. These species were identified through a multi-study, integrative analysis on 4,347 human stool metagenomes from 34 published studies across healthy and 12 different disease or abnormal bodyweight conditions. When demonstrated on our population-scale meta-dataset, GMHI is the most robust and consistent predictor of general health compared to α-diversity indices commonly considered as markers for gut health. Validation of GMHI on 679 samples from 9 additional studies resulted in remarkable reproducibility in distinguishing healthy and non-healthy groups. Our findings suggest that gut taxonomic signatures can indeed serve as robust predictors of general health, and highlight the importance of how data sharing efforts can provide broadly-applicable novel discoveries.

## INTRODUCTION

Recent advances in the field of human gut microbiome research have revealed novel associations and potential mechanistic insights regarding a vast array of complex, chronic diseases, including cancer^1,2^, autoimmune disease^3–5^, and metabolic syndrome^6–8^. Undoubtedly, the many microbiome studies that focused on disease contexts have been essential for elucidating underlying pathophysiological mechanisms, and for developing potential intervention strategies. However, in comparison, rigorous investigations into characterizing the general features of a healthy gut microbiome are considerably lacking, despite previous landmark efforts^9,10^. In this particular context, the search for an individual microbial species or strain that can promote health and wellness has been largely elusive; rather, mounting evidence suggests that the inter-relationship between a gut ecosystem and health is determined by the complex and collective properties of an ecological community^11–15^. Therefore, determining a particular set of gut commensals, and then understanding how they interact and function in unison, will be fundamental towards uncovering beneficial microbes that could maintain or restore health.

As researchers uncover more details regarding which gut commensals may play a significant role in host health, a promising application for this knowledge is towards developing a stool-based test to evaluate the health status tied to one’s own gut microbiome. Among several measures of gut ecological properties, Shannon diversity in particular has frequently been applied for comparing gut microbiomes implicated in health and disease^16–18^. So far, however, there is no conclusive evidence that such gross characteristics are a reliable and accurate measure to explain variations in health traits^19,20^. Therefore, the development of a simple and biologically interpretable metric that can quantify the degree of wellness (or the divergence away from a healthy condition) based on a gut microbiome snapshot will not only provide a significant advancement in current microbiome science, but also address a pressing clinical need. We can imagine that one can, through continuous monitoring, be able to detect significant changes or abnormalities in comparison to her/his normal baseline measurements; in turn, this will be the cue for additional diagnostic procedures and/or therapeutic interventions. Collecting diverse longitudinal biomolecular data from human subjects, and translating the corresponding complex datasets into actionable possibilities, has been previously demonstrated by Price *et al.*^21^

Since new insights regarding the gut microbiome and human health need to be reliable and robust across a wide range of human subjects and conditions, ‘population-level’ analyses can serve as an important platform for the discovery of broadly applicable principles and methodologies^22–26^. Traditionally, the collection of a substantial amount of microbiome samples (on the order of hundreds to thousands) for large-scale investigations has been undertaken at well-funded, major research centers and consortiums, mainly due to the prohibitive costs for lone laboratories. However, with the recent progress in the call for data sharing policies and practices^27^, conducting meta-analyses by crowdsourcing and/or re-purposing data from existing published studies have already begun to play important roles in either hypothesis generation or validation in microbiome research^1,28–31^. As 16s rRNA amplicon and shotgun metagenomic sequencing microbiome data are now readily available from multiple, independent studies conducted around the world, the meta-analysis approach provides a promising strategy to find health- and disease-associated signatures, as well as to gain new, holistic insights not offered by smaller, individual studies.

In this study, we introduce the Gut Microbiome Health Index (GMHI), a robust index for evaluating health status based on the species-level taxonomic profile of a stool shotgun metagenome (gut microbiome) sample. GMHI is designed to evaluate the balance between two sets of microbial species associated with good and adverse health conditions, which are identified from a meta-analysis on 4,347 publicly available, human stool metagenomes integrated across multiple studies encompassing various phenotypes and geographies. Applying GMHI to each sample in our population-scale meta-dataset, we find that GMHI distinguishes healthy from non-healthy groups far better than well-recognized ecological indices, e.g., Shannon diversity and richness. In addition, intra-study comparisons of stool metagenomes between healthy and non-healthy phenotypes demonstrate that GMHI is the most robust and consistent predictor of health. Finally, to confirm that GMHI classification performance was not a result of overfitting on the training data, we validate our approach on a previously unseen dataset of 679 samples from eight additional published studies and one new cohort. We find that GMHI demonstrates strong reproducibility in stratifying healthy and non-healthy groups, and furthermore, outperforms α-diversity indices generally considered as markers for gut health or dysbiosis.

## RESULTS

### A meta-dataset of human stool metagenomes integrated across 34 independent published studies

An overview of our multi-study integration approach, wherein we acquired 4,347 raw shotgun stool metagenomes (2,636 and 1,711 metagenomes from healthy and non-healthy individuals, respectively) from 34 independent published studies, is depicted in **Fig. 1**. In this study, ‘healthy’ subjects were defined as those who were reported as not having any overt disease nor adverse symptoms at the time of the original study; alternatively, ‘non-healthy’ subjects were defined as those who were clinically diagnosed with a disease, or determined to have abnormal bodyweight based on body mass index (BMI). Accordingly, 1,711 stool metagenomes from patients across 12 different disease or abnormal bodyweight conditions were pooled together into a single aggregate non-healthy group. (Our sample selection criteria are described in **Methods**. Importantly, all metagenomes were re-processed uniformly, thereby removing a major non-biological source of variance among different studies, as previously demonstrated^32^.) Descriptions of the studies whose human stool metagenomes were collected and processed through our study pipeline is provided in **Table 1**. We note that, in order to eventually identify features of the gut microbiome associated exclusively with health, it is important to be disease-agnostic by considering a broad range of non-healthy phenotypes. We provide all subjects’ phenotype, age, sex, BMI, and other questionnaire measures (as provided in their respective original study) in **Supplementary Table 1**. Along with the additional 679 stool metagenome samples used for validation purposes (discussed below), this study provides the largest metagenome meta-analysis of the human gut microbiome to date, in regards to the number of samples, phenotypes, and independent studies.

**Table 1.**
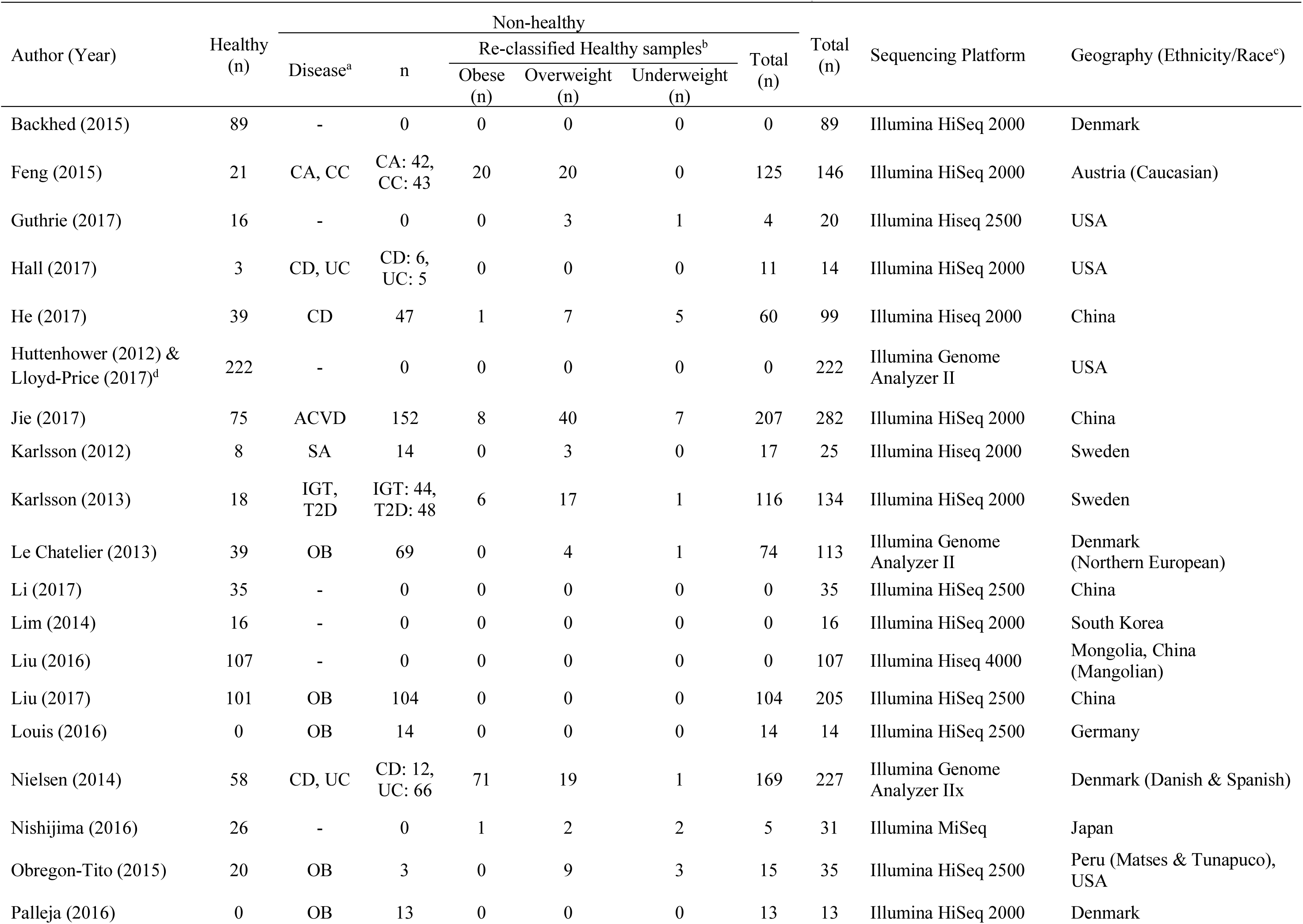
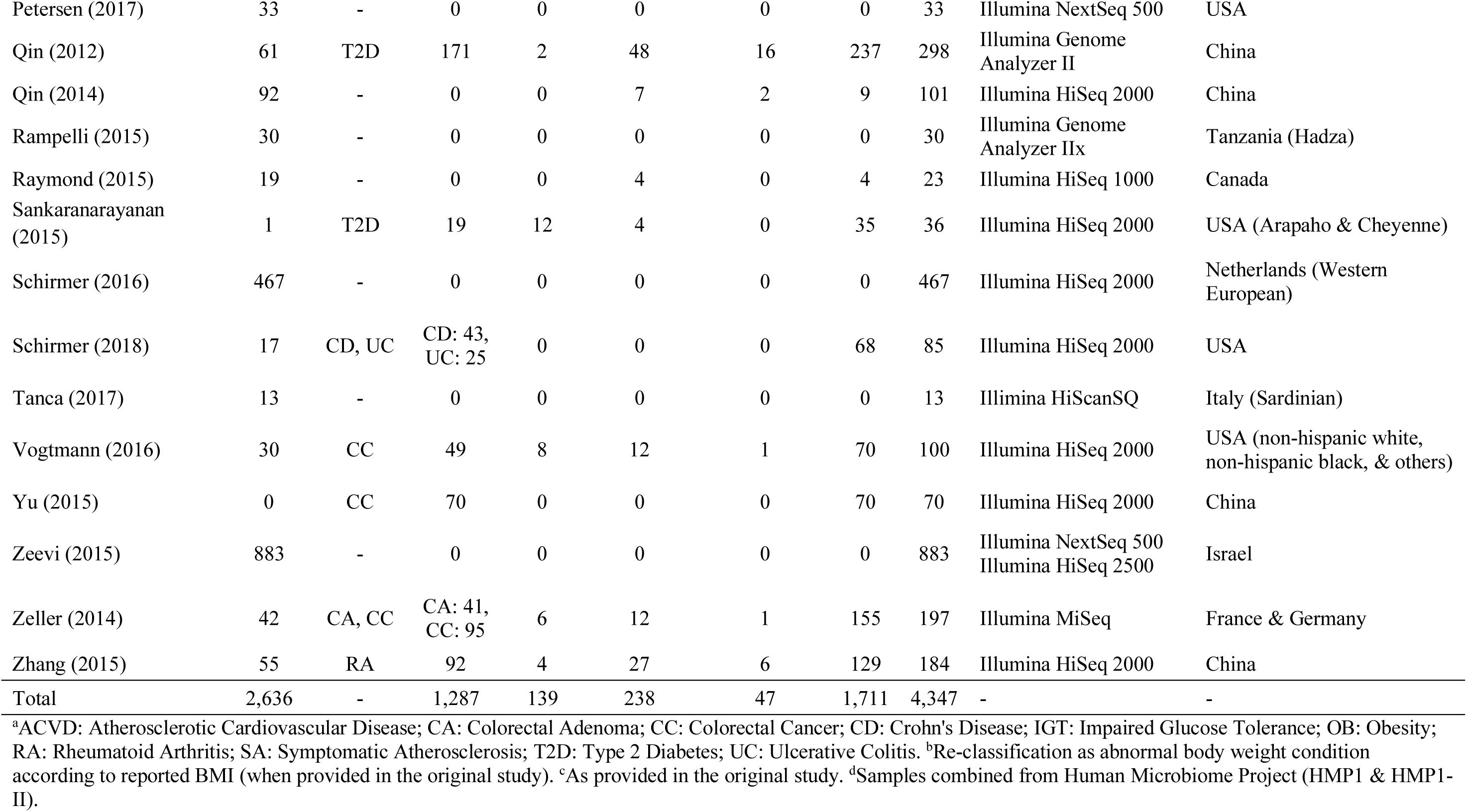
Human stool metagenome datasets collected, processed, and analyzed in this study.

**Figure 1.**
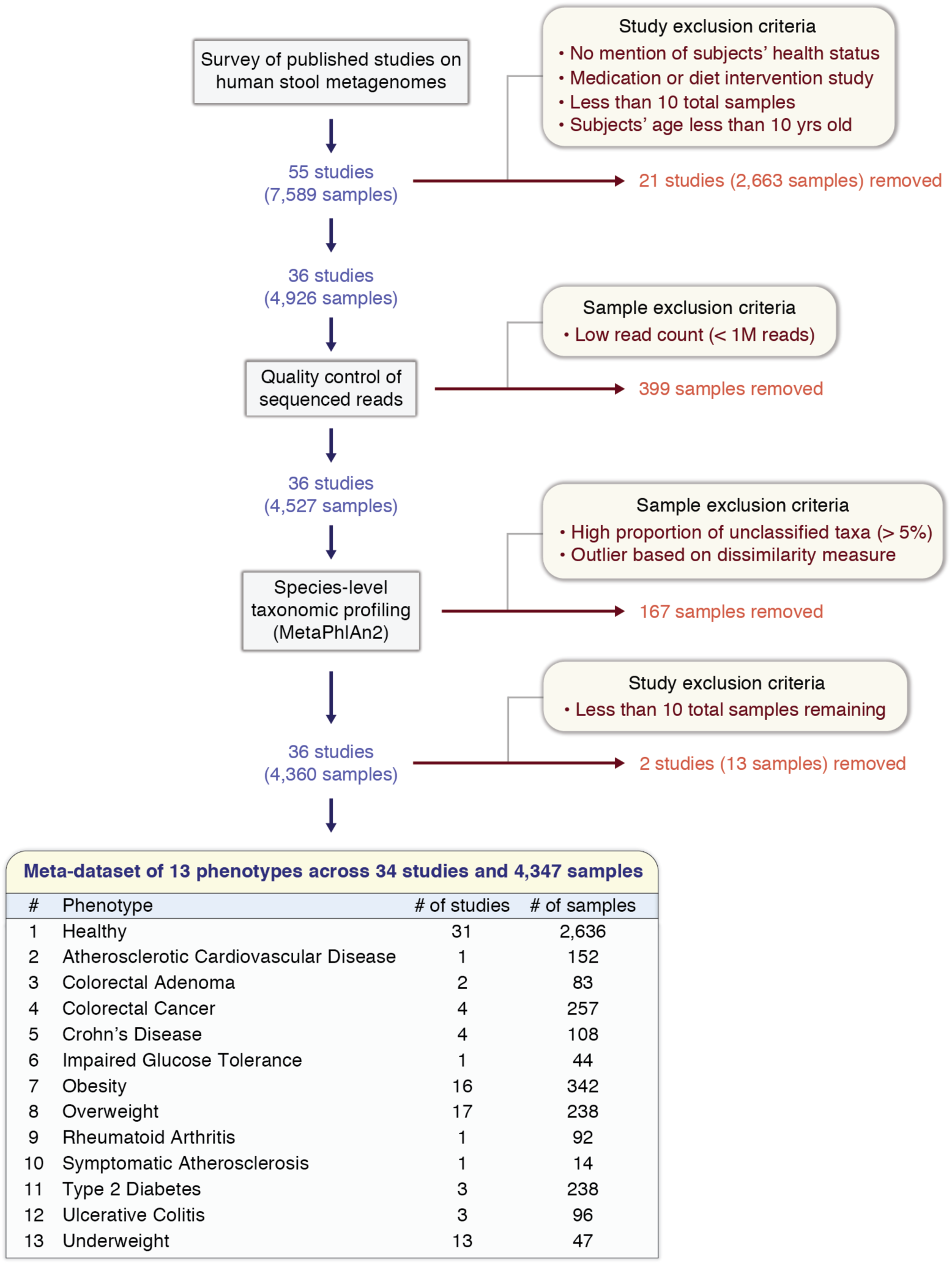
Schematic overview of the construction of a meta-dataset of human stool metagenomes integrated across multiple studies. A survey was conducted in PubMed and Google Scholar to search for published studies with publicly available human stool metagenome (gut microbiome) samples from healthy and non-healthy individuals. The initial collection of stool metagenomes consisted of 7,589 samples from 55 independent studies. All samples (.fastq files) were downloaded and re-processed uniformly using identical bioinformatics methods (see **Methods**). After quality control of sequenced reads, species-level taxonomic profiling was then performed. Studies and metagenome samples were removed based on several exclusion criterias. Finally, a total of 4,347 samples (2,636 and 1,711 metagenomes from healthy and non-healthy individuals, respectively) from 34 studies ranging across healthy and 12 non-healthy phenotypes were assembled into a meta-dataset for downstream analyses.

Importantly, we chose to integrate datasets from independent studies for two notable advantages: i) the expansion of sample number may help to amplify the primary biological signal of interest and improve statistical power^33,34^; and ii) the identified health/disease-associated signatures may encompass a wide range of heterogeneity across different sources and conditions (e.g., host genetics, geography, dietary and lifestyle patterns, age, sex, birth mode, early life exposures, medication history, stool consistency, sequencing depth), thereby helping to identify robust findings despite systematic biases from batch effects or other confounding factors^32,35^.

### Highly-prevalent ‘core’ members of the human gut microbiome

After downloading, re-processing, and performing quality filtration on all raw metagenomes, species-level taxonomic profiling was carried out using the MetaPhlAn2 pipeline^36^ (**Methods**). A total of 1,201 species were detected in at least one metagenome sample; after removing viruses, and species that were rarely observed or of unknown/unclassified identity (**Methods**), 313 species remained for further analysis (**Fig. 2a** and **Supplementary Table 2**; a phylogenetic tree showing the evolutionary relationships among these species is shown in **Supplementary Fig. 1**). Interestingly, six species (*Bacteroides ovatus, Bacteroides uniformis, Bacteroides vulgatus, Faecalibacterium prausnitzii, Ruminococcus obeum*, and *Ruminococcus torques*) were of high prevalence (i.e., detected in over 90% of all 4,347 samples), suggesting species-level members of a ‘core’ human gut microbiome.

**Figure 2.**
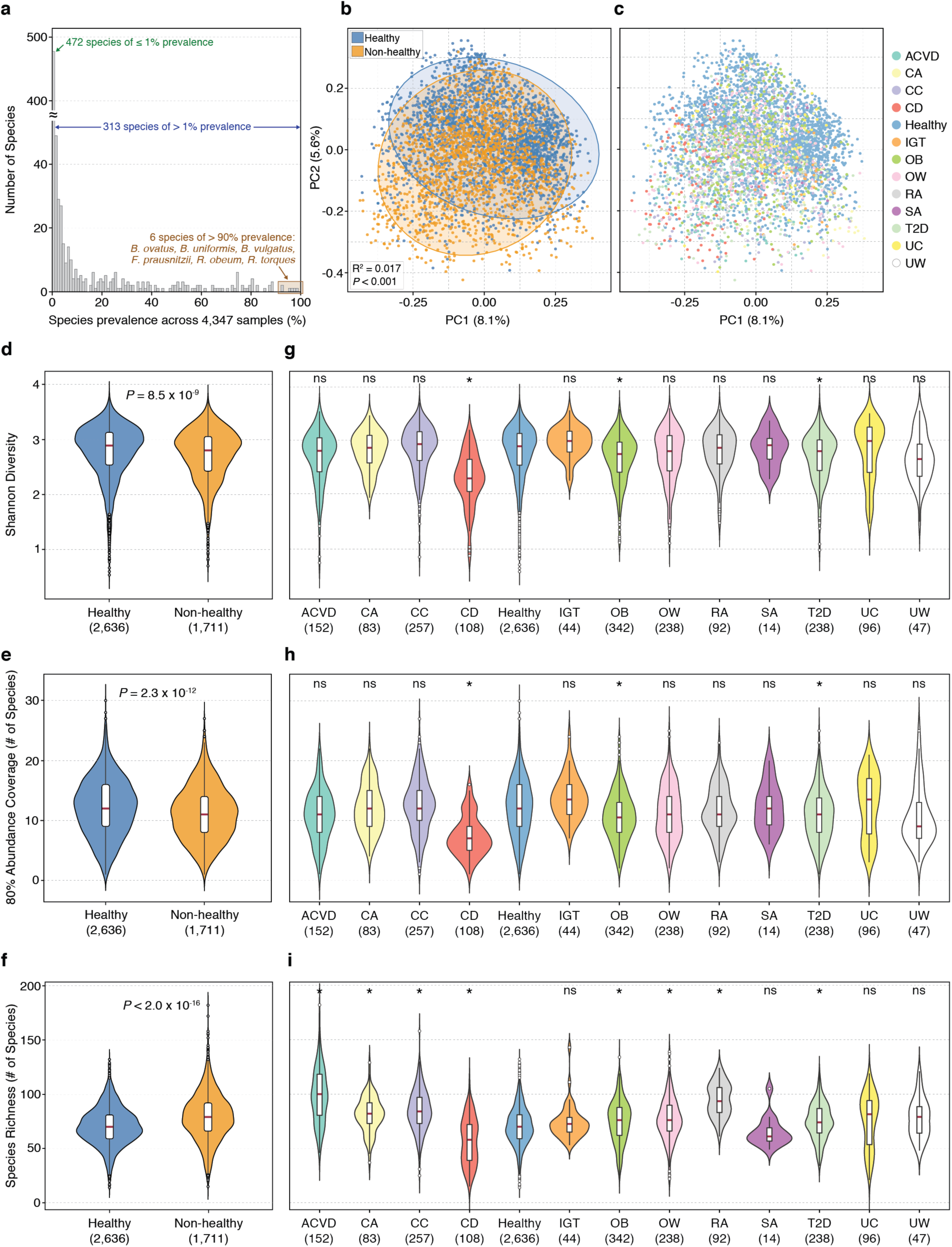
Healthy and non-healthy gut microbiomes show distinct ecological characteristics. **(a)** Distribution of microbial species’ prevalence across the 4,347 stool metagenome samples in the meta-dataset. After removing viruses, unknown/unclassified species-level entities, and rarely-observed species (i.e., detected < 1% of all samples), 313 species remained for further analyses. **(b)** Principal Coordinates Analysis (PCoA) ordination plot based on Bray-Curtis distances shows that healthy (blue; n = 2,636) and non-healthy (orange; n = 1,711) groups have significantly different distributions of gut microbiome profiles according to PERMANOVA (R^2^ = 0.017, *P* < 0.001). Each point corresponds to a sample. Ellipses correspond to 95% confidence regions. **(c)** In an identical PCoA plot, each color represents one of the thirteen different phenotypes of health or disease. **(d–f)** Stool metagenomes in healthy and non-healthy groups show significant differences in Shannon diversity, 80% abundance coverage, and species richness. Higher distributions of Shannon diversity and 80% abundance coverage were observed in gut microbiomes of healthy than in those of non-healthy individuals, whereas higher species richness was observed in non-healthy gut microbiomes. **(g–i)** Significant differences in Shannon diversity, 80% abundance coverage, and species richness were observed in comparisons between healthy and each of the twelve non-healthy phenotypes. For Shannon diversity and 80% abundance coverage, only three non-healthy phenotypes (CD, OB, and T2D) were found to have different distributions compared to healthy; both properties were higher in healthy than in CD, OB, and T2D. For species richness, seven (ACVD, CA, CC, OB, OW, RA, and T2D) of the twelve non-healthy phenotypes were observed to have significantly higher richness than healthy; in contrast, only CD showed significantly lower richness compared to healthy. *P*-values shown above the violin plots were found using Mann-Whitney *U* test. *, *P* < 0.001 in Mann-Whitney *U* test; **ns**, not significant. The sample size of each group is shown in parentheses. ACVD, atherosclerotic cardiovascular disease; CA, colorectal adenoma; CC, colorectal cancer; CD, Crohn’s disease; IGT, impaired glucose tolerance; OB, obesity; OW, overweight; RA, rheumatoid arthritis; SA, symptomatic atherosclerosis; T2D, type 2 diabetes; UC, ulcerative colitis; and UW, underweight.

### Species-level analyses identify significant differences between healthy and non-healthy gut ecologies

The overall ecology of the gut microbiome has often been associated with host health^8,37–39^. Using species-level relative abundance (i.e., proportion) profiles, we examined for differences in gut microbial diversity between the healthy (n = 2,636) and non-healthy (n = 1,711) groups. First, when using Principal Coordinates Analysis (PCoA) ordination (on a Bray-Curtis distance matrix), we identified a significant difference between the distributions of these two groups (R^2^ = 0.017, *P* < 0.001, PERMANOVA; and **Fig. 2b**). In the same PCoA plot in which the healthy and twelve non-healthy phenotypes were presented simultaneously, we found that the various phenotypes were homogeneously dispersed without any apparent sub-clusters (**Fig. 2c**).

Next, to further identify differences between healthy and non-healthy groups, we examined multiple measures of ecological characteristics that can be defined on a per-sample basis. For α-diversity based on the Shannon index, we found significantly higher values in healthy than in non-healthy (*P* = 8.5×10^−9^, Mann-Whitney *U* test; and **Fig. 2d**). In agreement with our results, previous investigations have also reported higher diversity in the gut microbiomes of healthy controls than in those of disease patients^40–42^; however, these studies consist of specific case-control comparisons and were conducted on much smaller sample sizes than our current study. In addition, we found that the minimum number of species required to comprise at least 80% of the sample’s relative abundance (henceforth called ‘80% abundance coverage’) was significantly higher in healthy compared to non-healthy (*P* = 2.3×10^−12^, Mann-Whitney *U* test; and **Fig. 2e**). This concept, as demonstrated similarly by Kraal *et al*.^43^, has been adopted in previous studies to estimate the membership of core microbiomes^44,45^. Finally, species richness, which is defined as the observed number of different species, was found to be significantly lower in healthy compared to non-healthy (*P* < 2.0×10^−16^, Mann-Whitney *U* test; and **Fig. 2f**).

Finally, we compared these ecological characteristics between healthy and each of the twelve phenotypes of the non-healthy group. For Shannon diversity and 80% abundance coverage, we found that only three (Crohn’s Disease, Obesity, and Type 2 diabetes) of the twelve non-healthy phenotypes showed statistically significant differences (**Figs. 2g** and **2h**); both properties were higher in healthy for all three comparisons. For richness, we found that eight of the twelve non-healthy phenotypes were significantly different compared to healthy (**Fig. 2i**): seven of these eight were of higher richness, whereas the remaining one (Crohn’s Disease) was of lower richness. In summary, our analysis shows that healthy and non-healthy gut microbiomes show distinct ecological characteristics, although no consistent and reliable trend explaining the variability was observed for any of these measures in phenotype-level comparisons.

### A prevalence-based strategy for identifying microbial species associated with healthy human gut microbiomes

We set out to identify distinct microbial species associated with healthy (*H*) and non-healthy (*N*) groups. Here, we used a prevalence-based strategy to deal with the sparse nature of microbiome datasets. For this, we first determine 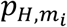 and 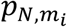, or the prevalence of microbial species *m*_*i*_, i.e., proportion of samples in a given group wherein *m*_*i*_ is ‘present’ (or relative abundance ≥ 1.0×10^−5^) in *H* and *N*, respectively. Next, for comparing the two prevalences in *H* and *N*, we apply the following two criteria: prevalence fold-change 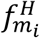 and prevalence difference 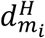, defined as 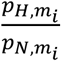 and 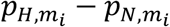, respectively.

A significant effect size between the two prevalences is considered to exist if both criteria satisfy (predetermined) minimum thresholds for prevalence fold-change *θ*_*f*_ and prevalence difference *θ*_*d*_ (how we determine the best pair of thresholds for distinguishing *H* and *N* is described below). For all detectable microbial species that satisfy 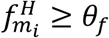 and 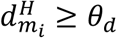, we term these species that are more frequently observed in *H* (than in *N*) as ‘Health-prevalent’ species *M*_*H*_. Analogously, we identify ‘Health-scarce’ species *M*_*N*_, or the species that are less frequently observed in *H* (than in *N*), as those that satisfy 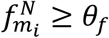 and 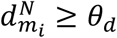, where 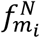 and 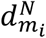 for *m*_*i*_ are defined as 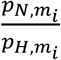 and 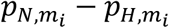, respectively. In this regard, the species that are eventually chosen to compose *M*_*H*_ and *M*_*N*_ both dependent on *θ*_*f*_ and *θ*_*d*_. An important strength of our prevalence-based strategy for identifying microbial associations is that it does not compare ensemble-averaged measurements between the two groups, which is challenging to justify when biological and technical heterogeneity (batch effects) could vary greatly across various independent studies. Rather, our approach compares frequencies of a ‘present’ signal—on a sample-by-sample basis—between two groups, and may offer a more robust strategy applicable to the context of integrating high-throughput data across different studies.

### Balancing the collective abundances of two sets of microbial taxonomies

Having a strategy to identify microbial species associated with healthy (i.e., Health-prevalent or *M*_*H*_) and non-healthy (i.e., Health-scarce or *M*_*N*_), we next couple these two sets of taxonomies with a computational procedure that quantifies the health status of any gut microbiome sample. To this end, we developed the following mathematical formula: for a given set *M*_*H*_ (Health-prevalent species) in sample *i*, their ‘collective abundance’ 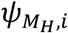 is defined as

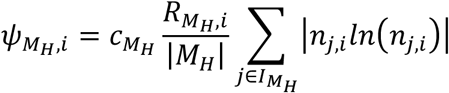

where 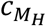 is a constant specific to *M*_*H*_, 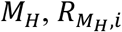 is the richness of species in *M*_*H*_ in sample *i*, | *M*_*H*_| is the set size of *M*_*H*_, 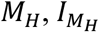 is the index set of *M*_*H*_,, and *n*_*j,i*_ is the relative abundance of species *j* in sample *i*. Full details on the physical meaning and derivation of 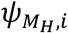 are described in **Supplementary Methods**. For the microbial species in another given set *M*_*N*_ (Health-scarce species) in the same sample *i*, their ‘collective abundance’ 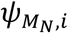 can be defined analogously. Next, the collective abundances of species in sets *M*_*H*_ and *M*_*N*_ in sample *i* are compared by identifying the ratio of 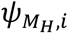 to 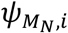 as

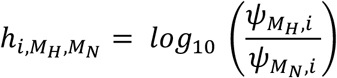

where 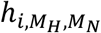 denotes the degree to which sample *i* portrays the collective abundance of either set *M*_*H*_ or set *M*_*N*_,. More specifically, a positive or negative 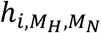 suggests that sample *i* is characterized more by the microbial properties of *M*_*H*_ or *M*_*N*_, respectively; an 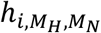 of 0 indicates that there is an equal balance of both traits. Full details on the derivation of 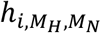 are provided in **Supplementary Methods**.

### Determining optimal sets of Health-prevalent and Health-scarce species

As introduced above, we use minimum thresholds *θ*_*f*_ and *θ*_*d*_ for prevalence fold-change and prevalence difference to control for the number of Health-prevalent and Health-scarce species; species that simultaneously satisfy the two types of thresholds are selected to be included in one of either group. Next, Health-prevalent species (assigned as set *M*_*H*_) and Health-scarce species (assigned as set *M*_*N*_) can be provided as input features for 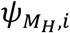 and 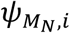, respectively, and for the calculation of 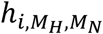, which in turn can classify stool metagenome sample *i* as healthy (i.e., 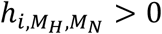), non-healthy (i.e., 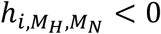), or neither (i.e., 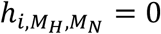). Lastly, we test 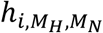 on all 4,347 stool metagenomes in our meta-dataset by finding the average classification accuracy 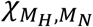, defined as the average of the proportions of healthy and non-healthy samples that were correctly classified, or

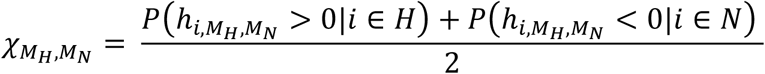

where 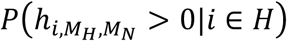 is the proportion of samples in the healthy group (*H*) whose 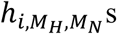 are positive, and 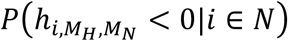 is the proportion of samples in the non-healthy group (*N*) whose 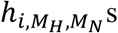 s are negative.

We determine the final, optimal sets of Health-prevalent and Health-scarce species (and their corresponding prevalence thresholds) that result in the highest average classification accuracy 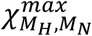. After systematically testing across a range of two different thresholds (every pair of *θ*_*f*_ and *θ*_*d*_ gives different sets of *M*_*H*_ and *M*_*N*_, and in turn a different 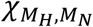), we observed 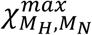 when *θ*_*f*_ and *θ*_*d*_ were set to 1.4 and 10%, respectively (**Supplementary Table 3**). We identified 50 microbial species that satisfy both of these thresholds simultaneously; among these 50 species, 7 and 43 comprise the Health-prevalent and Health-scarce groups, respectively (**Table 2**). We were able to confirm higher relative abundance levels of Health-prevalent and Health-scarce species in the healthy and non-healthy group, respectively (**Supplementary Fig. 2**). Furthermore, in **Supplementary Fig. 3**, we show the prevalence of these species in case (i.e., non-healthy) and/or control (i.e., healthy) for the 34 published studies upon which our stool metagenome meta-dataset was derived. Despite the heterogeneity and unevenness in prevalences across all studies, we found that, by and large, Health-prevalent and Health-scarce species were observed more frequently in the control and case samples, respectively. Finally, we report known associations between Health-prevalent/-scarce species and health/disease in **Supplementary Table 4**.

**Table 2.**
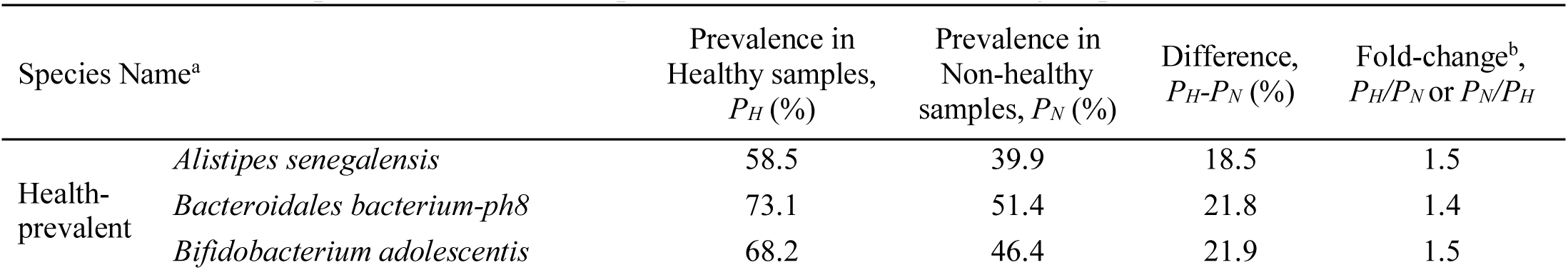
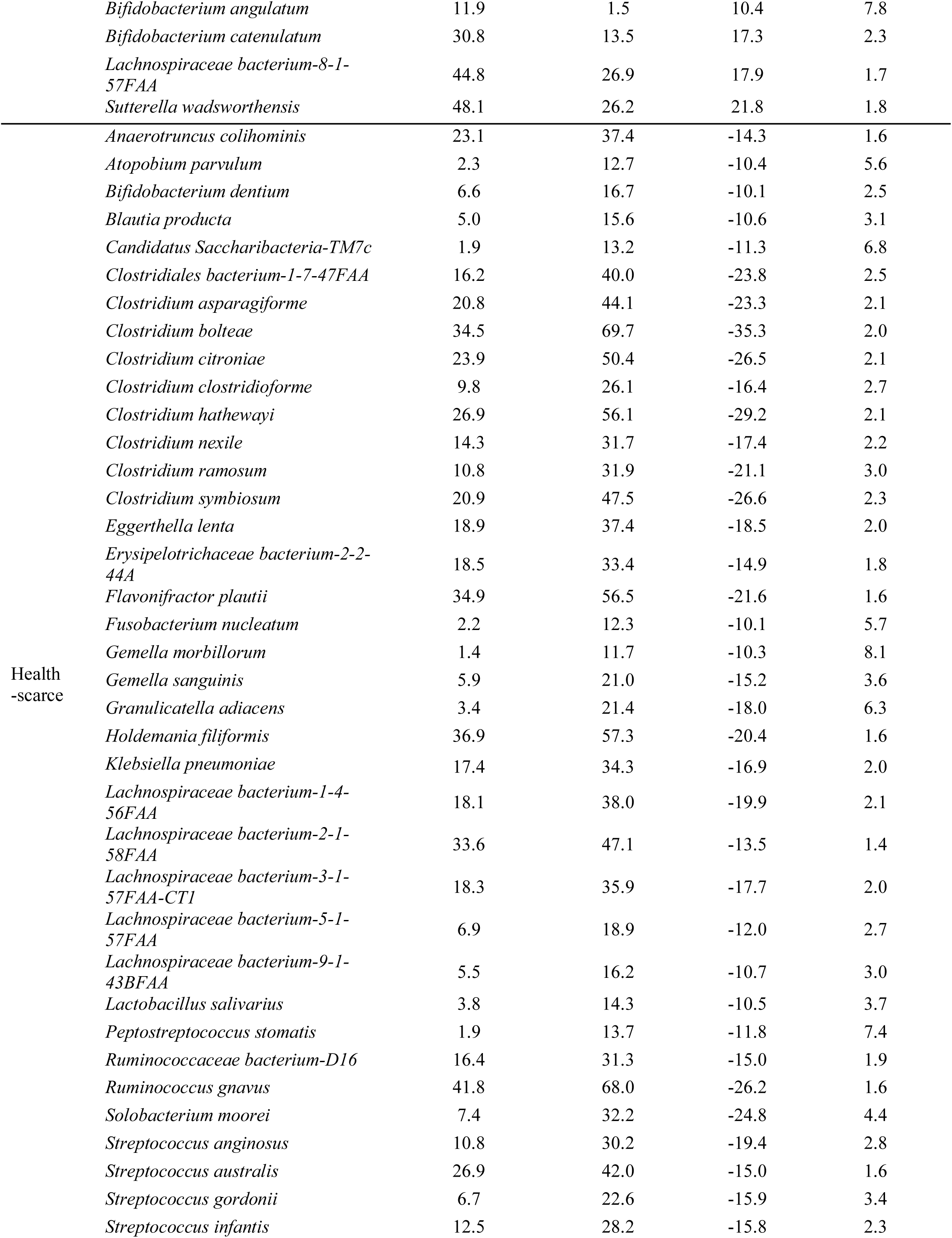
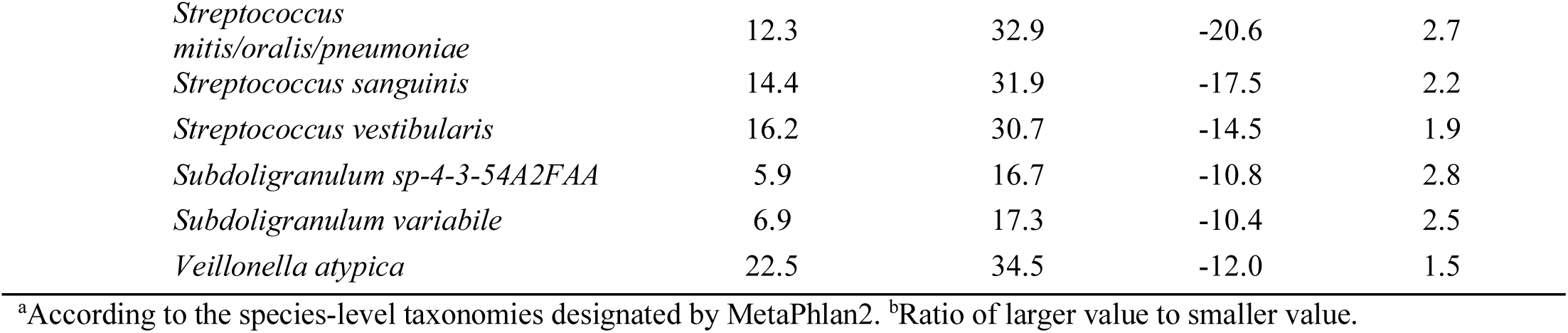
Microbial species of the Health-prevalent and Health-scarce groups.

### Gut Microbiome Health Index: A metric for predicting health status using gut microbiome

Henceforth, we term the ratio 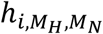 between these two groups of 7 Health-prevalent and 43 Health-scarce species as the Gut Microbiome Health Index (GMHI). GMHI accumulates the count of Health-prevalent and Health-scarce species observed to be present in a microbiome sample, their relative abundances, and their diversity (based on Shannon index), into one quantitative metric. Collectively, GMHI denotes the degree to which a subject’s stool metagenome sample portrays microbial taxonomic properties associated with healthy or non-healthy.

Analogous to the example mentioned above, a positive or negative GMHI allows the sample to be classified as healthy or non-healthy, respectively; a GMHI of 0 indicates an equal balance of Health-prevalent and Health-scarce species, and thereby classified as neither. Therefore, GMHI is especially favorable in terms of the simplicity of the decision rule and the biological interpretation regarding the two sets of microbes involved in classification. As GMHI is measured on an individual microbiome sample, it could be used not only to measure the health (or dysbiotic) status of one’s gut microbial community, but also to assess the deviation of one’s gut microbiome from a presumed healthy state. Importantly, our metric requires very little parameter-tuning and foregoes the use of arbitrary thresholds, e.g., ‘low’ or ‘high’ α-diversity.

### Gene copy number variation analysis identifies potential metabolic functions specific to Health-prevalent species

To uncover genes that could explain differences in functional potential between the Health-prevalent (n = 7) and Health-scarce (n = 43) species, we constructed gene (KEGG ortholog) copy number profiles of all 50 species using genomes of reference strains (**Methods** and **Supplementary Table 5**). In **Fig. 3**, we show the 62 KEGG functional orthologs that were identified to have significant differences in copy number between the Health-prevalent and Health-scarce species (*q* < 0.01, Mann-Whitney *U* test with Benjamini-Hochberg FDR correction; **Supplementary Table 6**). Hierarchical clustering revealed that five of the seven Health-prevalent microbes (*Alistipes senegalensis, Bacteroides bacterium-ph8, Bifidobacterium adolescentis, Bifidobacterium angulatum*, and *Bifidobacterium catenulatum*) tightly clustered based on their gene copy number profiles, especially by enzymes involved in the metabolism of sugars and polysaccharides (α-galactosidase, β-galactosidase, β-glucosidase, glycogen debranching enzyme, maltose O-acetyltransferase, L-rhamnosyltransferase, and sucrose-6-phosphatase) and lipids (long-chain acyl-CoA synthetase).

**Figure 3.**
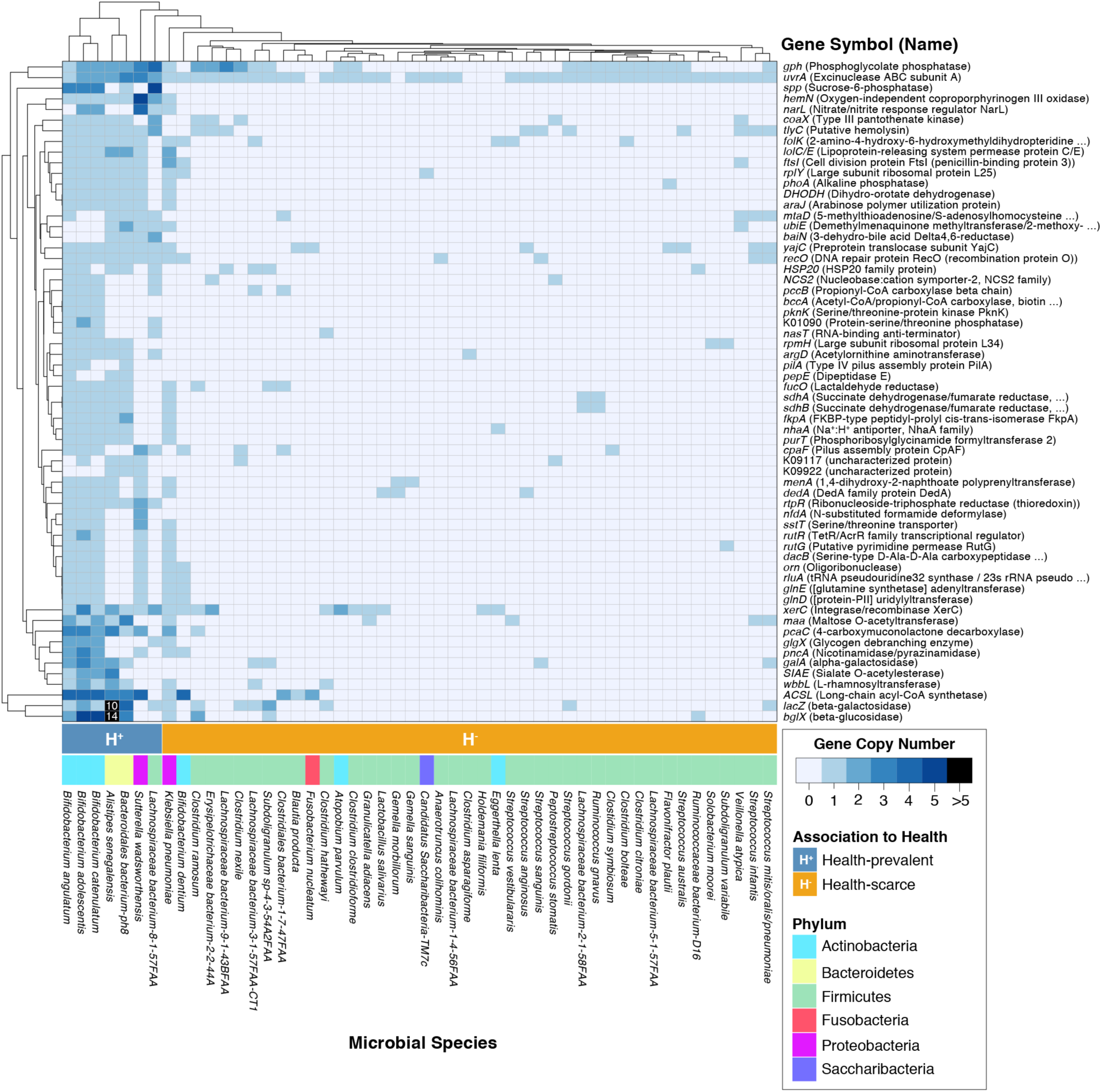
Analysis of gene copy number variation on reference strains reveals KEGG functional orthologs significantly increased in microbial species prevalent in healthy. 62 KEGG functional orthologs were identified to have significant variation in copy number between reference genomes of the Health-prevalent (n = 7) and Health-scarce (n = 43) species (*q* < 0.01). All genes with annotated functional orthologs in KEGG were considered for this analysis, but only those found to have significant gene copy number variation between the two groups of species are shown. On the left side of the heatmap, five (*A. senegalensis, B. adolescentis, B. angulatum, B. bacterium-ph8*, and *B. catenulatum*) of the seven Health-prevalent species tightly clustered together while the remaining two species (*L. bacterium* 8-1-57FAA and *S. wadsworthensis*) situated adjacent to this cluster. Health-scarce species clustered into several sub-groups. All 62 genes were found to have significantly higher copy numbers in Health-prevalent species than in Health-scarce species. Mann-Whitney *U* test was used to calculate *P*-values, and Benjamini-Hochberg approach was used to control for false discovery rate. Full names of genes are available in **Supplementary Table 6**.

Interestingly, all 62 genes were found to have higher copy numbers in Health-prevalent species than in Health-scarce species. The detection of such genes, which include many metabolic pathway enzymes, offers insight into the underlying functional roles of microbes prevalent in the gut of healthy individuals. Additionally, through KEGG pathway enrichment analysis (**Supplementary Table 7**), we found that these genes with differential copy number were over-represented in metabolic pathways including ‘Sphingolipid Metabolism’ (*P* = 0.016, hypergeometric test) and ‘Glycosphingolipid Biosynthesis’ (*P* = 0.045, hypergeometric test). A recent study in mice by Brown *et al.* showed that the deficiency in the production of bacterial sphingolipids correlates with inflammatory bowel disease severity, and that bacterial sphingolipids help to maintain intestinal homeostasis and microbial symbiosis in host gut^46^. Although it remains unclear regarding how, and in which context, these complex lipids are produced or metabolized by Health-prevalent species, our findings nevertheless shed further light on possible health implications of gut microbiome-derived sphingolipids.

### GMHI stratifies healthy and non-healthy groups using species-level abundance profiles

We calculated GMHI for each stool metagenome in our meta-dataset of 4,347 samples to investigate whether the distributions of GMHI differ between the healthy and non-healthy groups. We found that the gut microbiomes in healthy have significantly higher GMHIs in comparison to gut microbiomes in non-healthy (*P* < 2.0×10^−16^, Mann-Whitney *U* test; **Fig. 4a**). By definition of GMHI, this result reflects the dominant influence of Health-prevalent species over Health-scarce species in the healthy group, and vice versa in the non-healthy group. In regards to classification performance, GMHI achieved a 71.0% (3,084 of 4,347) classification accuracy when applying GMHI onto the training data (4,347 stool metagenome samples) itself. Interestingly, for increasingly higher (more positive) and lower (more negative) values of GMHI, we observed an increasing proportion of samples from healthy and non-healthy, respectively (**Fig. 4b**). For example, 98.2% (165 of 168) of metagenome samples with GMHIs higher than 4.0 were from the healthy group; and 81.2% (164 of 202) of metagenome samples with GMHIs lower than −4.0 were of non-healthy origin. Interestingly, the top 25 to 100 healthy and non-healthy stool metagenome groups (selected based on their GMHI) clearly clustered apart from each other in PCoA ordination (**Supplementary Fig. 4**), in stark contrast to the case when all samples were projected simultaneously (**Fig. 2b**). These observations confirm that very high (or low) cumulative abundance of Health-prevalent species relative to that of Health-scarce species is strongly connected to the individual being healthy (or non-healthy).

**Figure 4.**
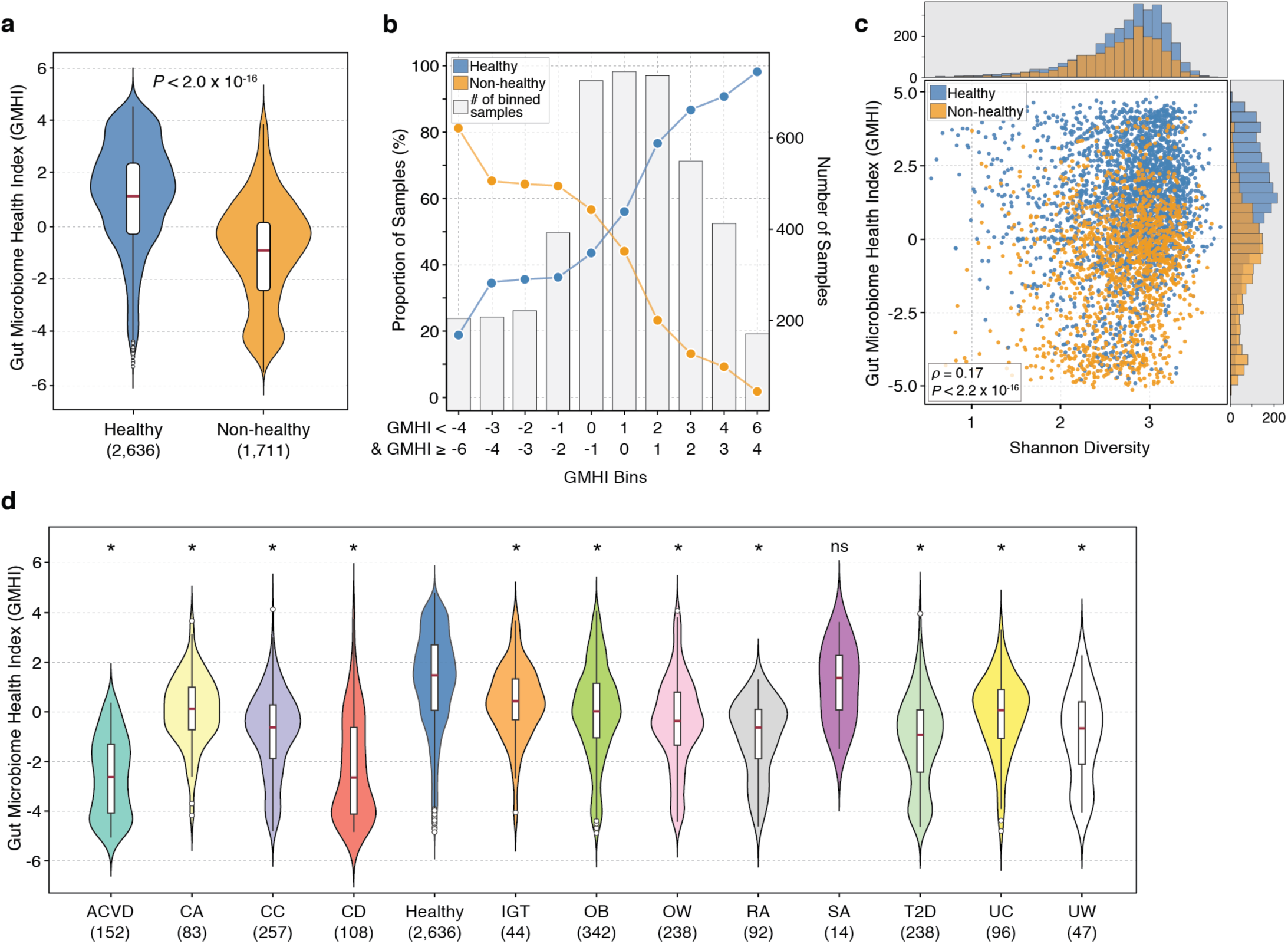
Predicting health status with Gut Microbiome Health Index (GMHI). **(a)** GMHIs from stool metagenomes of the healthy group are significantly higher than those of the non-healthy group. **(b**) All 4,347 metagenomes were binned according to their GMHI values (x-axis). Each bar (gray) indicates the total number of samples in each bin. Points indicate proportions (i.e., percentages) of samples in each bin corresponding to either healthy or non-healthy individuals (y-axis). In bins with a positive range of GMHIs, the majority of samples classified as healthy; in contrast, samples in bins with a negative range of GMHIs mostly classified as non-healthy. This trend was more pronounced towards bins on the far right and left. **(c**) GMHI stratifies healthy and non-healthy groups more strongly compared to Shannon diversity. Each point in the scatter-plot corresponds to a metagenome sample. Histograms show the distribution of healthy (blue) and non-healthy (orange) samples based on the parameter of each axis. In general, GMHI and Shannon diversity demonstrate a weak correlation (Spearman’s *ρ* = 0.17, *P* < 2.2×10^−16^). **(d**) The healthy group was found to have a significantly higher distribution of GMHIs than all but one (SA) of the twelve non-healthy phenotypes. In this regard, GMHI outperforms other ecological characteristics. *P*-value shown above the violin plot was found using Mann-Whitney *U* test. *, *P* < 0.001 in Mann-Whitney *U* test; **ns**, not significant. The sample size of each group is shown in parentheses. ACVD, atherosclerotic cardiovascular disease; CA, colorectal adenoma; CC, colorectal cancer; CD, Crohn’s disease; IGT, impaired glucose tolerance; OB, obesity; OW, overweight; RA, rheumatoid arthritis; SA, symptomatic atherosclerosis; T2D, type 2 diabetes; UC, ulcerative colitis; and UW, underweight.

### GMHI stratifies healthy and non-healthy groups more strongly than Shannon diversity

To examine their overall concordance, GMHI and Shannon diversity were compared for each sample in our meta-dataset. As shown in **Fig. 4c**, GMHI clearly performed much better in stratifying the healthy and non-healthy groups compared to Shannon diversity. Additionally, only a weak relationship was found between our metric and this conventional measure of gut health (Spearman’s *ρ* = 0.17, *P* < 2.2×10^−16^). Moreover, similar results were seen when GMHI was compared to 80% abundance coverage (Spearman’s *ρ* = 0.22, *P* < 2.2×10^−16^) and to richness (Spearman’s *ρ* = −0.25, *P* < 2.2×10^−16^) (**Supplementary Fig. 5**). At the individual phenotype-level, the healthy group showed significantly higher GMHI levels in all but one (symptomatic atherosclerosis) of the twelve different disease or abnormal bodyweight conditions (*P* < 0.001, Mann-Whitney *U* test; **Fig. 4d**). Taken together, our results suggest that GMHI: i) embodies a gut microbial signature of wellness that is generalizable against various non-healthy phenotypes; and ii) can distinguish healthy from non-healthy individuals more reliably than Shannon diversity, 80% abundance coverage, and richness (**Figs. 2d–i**).

### Intra-study comparisons of stool metagenome ecological characteristics between healthy and non-healthy phenotypes show GMHI as the most robust and consistent predictor of general health status

Having identified significant alterations between healthy and non-healthy groups when gut microbiomes were pooled together from various studies, we next examined how well GMHI and other features of microbial ecology (i.e., Shannon diversity, 80% abundance coverage, and species richness) could distinguish healthy and non-healthy phenotypes within individual studies. Specifically, in each of the fourteen studies (out of 34 total) wherein stool metagenome samples from both case (i.e., disease or abnormal bodyweight conditions) and control (i.e., healthy) subjects were available, we compared GMHI, Shannon diversity, 80% abundance coverage, and species richness between healthy and non-healthy phenotype(s). By focusing one-by-one on datasets from individual studies, this approach not only removes a major source of batch effects, but also provides a good means to investigate the robustness of our previously observed trends (when healthy and non-healthy samples were compared against each other in aggregate groups) across multiple, smaller studies.

We found that GMHI in healthy was significantly higher than that in any non-healthy phenotype for eleven out of thirty case-control comparisons (**Fig. 5a**). For Shannon diversity and 80% abundance coverage, we found significantly higher values in healthy than in non-healthy phenotypes for two and four case-control comparisons, respectively (**Figs. 5b** and **5c**). Lastly, we found species richness in healthy to be significantly lower than that in non-healthy phenotypes for five case-control comparisons (**Fig. 5d**). Clearly, the performance of GMHI was not perfect (and likewise for other ecological characteristics), as the expected trend was not replicable for all case-control comparisons within every study; overall though, GMHI strongly outperforms other microbiome ecological characteristics in distinguishing case and control.

**Figure 5.**
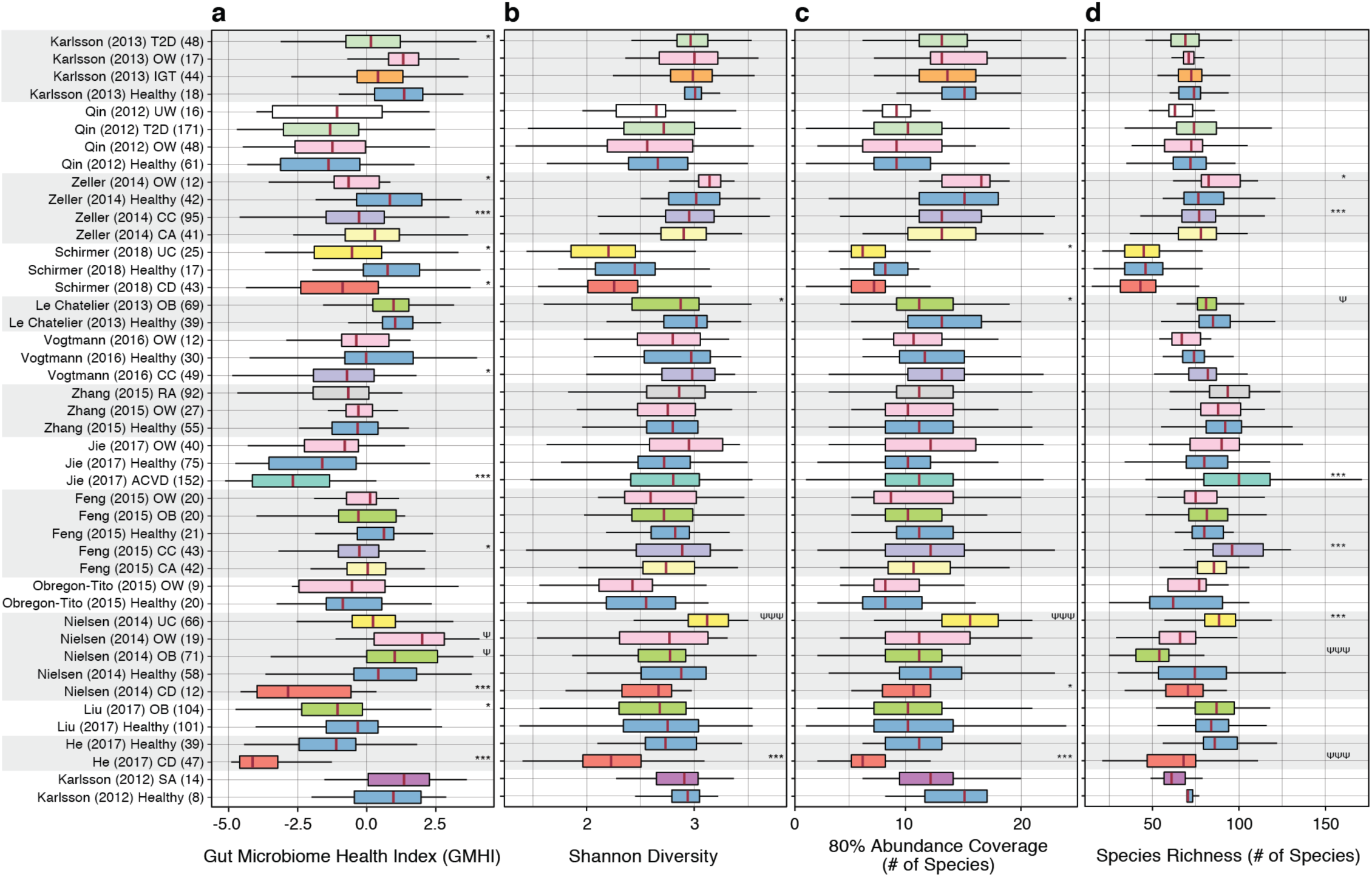
GMHI generally outperforms other microbiome ecological characteristics in distinguishing case and control across multiple study-specific comparisons. In each of the fourteen studies wherein case (i.e., disease or abnormal bodyweight conditions) and control (i.e., healthy) subjects were available, stool metagenomes were analyzed to compare **(a)** GMHI, **(b)** Shannon diversity, **(c)** 80% abundance coverage, and **(d)** species richness between healthy and non-healthy phenotype(s). GMHI was found to have a significantly higher distribution in healthy for eleven case-control comparisons across nine different studies; Shannon diversity and 80% abundance coverage were found to have significantly higher distributions in healthy for two and four case-control comparisons (across two and four studies), respectively; and species richness was found to have a significantly lower distribution in healthy for five case-control comparisons across four different studies. Each study’s phenotype sample size is shown in parentheses to the right of the phenotype abbreviation. The same colors in box-plots were used for the same phenotypes. *P*-values (Mann-Whitney *U* test) for each study-specific comparison between healthy and non-healthy phenotypes are shown adjacent to the boxplots. * and Ψ indicates significantly different distributions consistent with, and opposite to, respectively, the previously observed results when healthy and non-healthy groups were compared in aggregate. *** or ΨΨΨ, *P* < 0.001; * or Ψ, *P* < 0.05. ACVD, atherosclerotic cardiovascular disease; CA, colorectal adenoma; CC, colorectal cancer; CD, Crohn’s disease; IGT, impaired glucose tolerance; OB, obesity; OW, overweight; RA, rheumatoid arthritis; SA, symptomatic atherosclerosis; T2D, type 2 diabetes; UC, ulcerative colitis; and UW, underweight.

### Validation of GMHI on independent cohorts demonstrates strong reproducibility in stratifying healthy and non-healthy groups

Evaluation of any biomarker or molecular signature on fully independent patient samples is the gold standard for assessing its performance^47^. To confirm the reproducibility of our prediction results in stratifying healthy and non-healthy phenotypes (**Fig. 4**), we leveraged GMHI to predict the health status of 679 individuals whose stool metagenome samples were not part of the training dataset used for the original formulation of GMHI. For this, we used gut microbiome data from an additional eight published studies (**Supplementary Tables 8** and **9**), which include stool metagenomes from healthy subjects and patients with ankylosing spondylitis (AS), colorectal adenoma (CA), colorectal cancer (CC), Crohn’s disease (CD), liver cirrhosis (LC), and non-alcoholic fatty liver disease (NAFLD). In addition, we also collected our own cohort of patients with rheumatoid arthritis (RA) (**Methods**; see **Supplementary Table 9** for subject meta-data relating to both clinical and non-clinical factors). All metagenome samples in this validation dataset were pooled into one of two groups (i.e., healthy or non-healthy), as demonstrated above.

In agreement with our results on the training data, GMHIs from stool metagenomes of the healthy validation group (n = 118) were significantly higher than those of the non-healthy validation group (n = 561) (*P* < 2.0×10^−16^, Mann-Whitney *U* test; **Fig. 6a**). Next, we considered classification accuracy, which is defined simply as the proportion of healthy and non-healthy samples with positive and negative GMHIs, respectively. The classification accuracies for the healthy and non-healthy validation groups were 77.1% (91 of 118) and 70.2% (394 of 561), respectively. Notably, these accuracies were slightly better than the performances on the training dataset of 4,347 samples, which were 75.6% (1,993 of 2,636) and 63.8% (1,092 of 1,711) for healthy and non-healthy groups, respectively. Finally, we found that the proportions of accurate classifications for all samples of the training and validation datasets were nearly identical: 71.0% (3,084 of 4,347) and 71.4% (485 of 679) classification accuracy on training and validation datasets, respectively.

**Figure 6.**
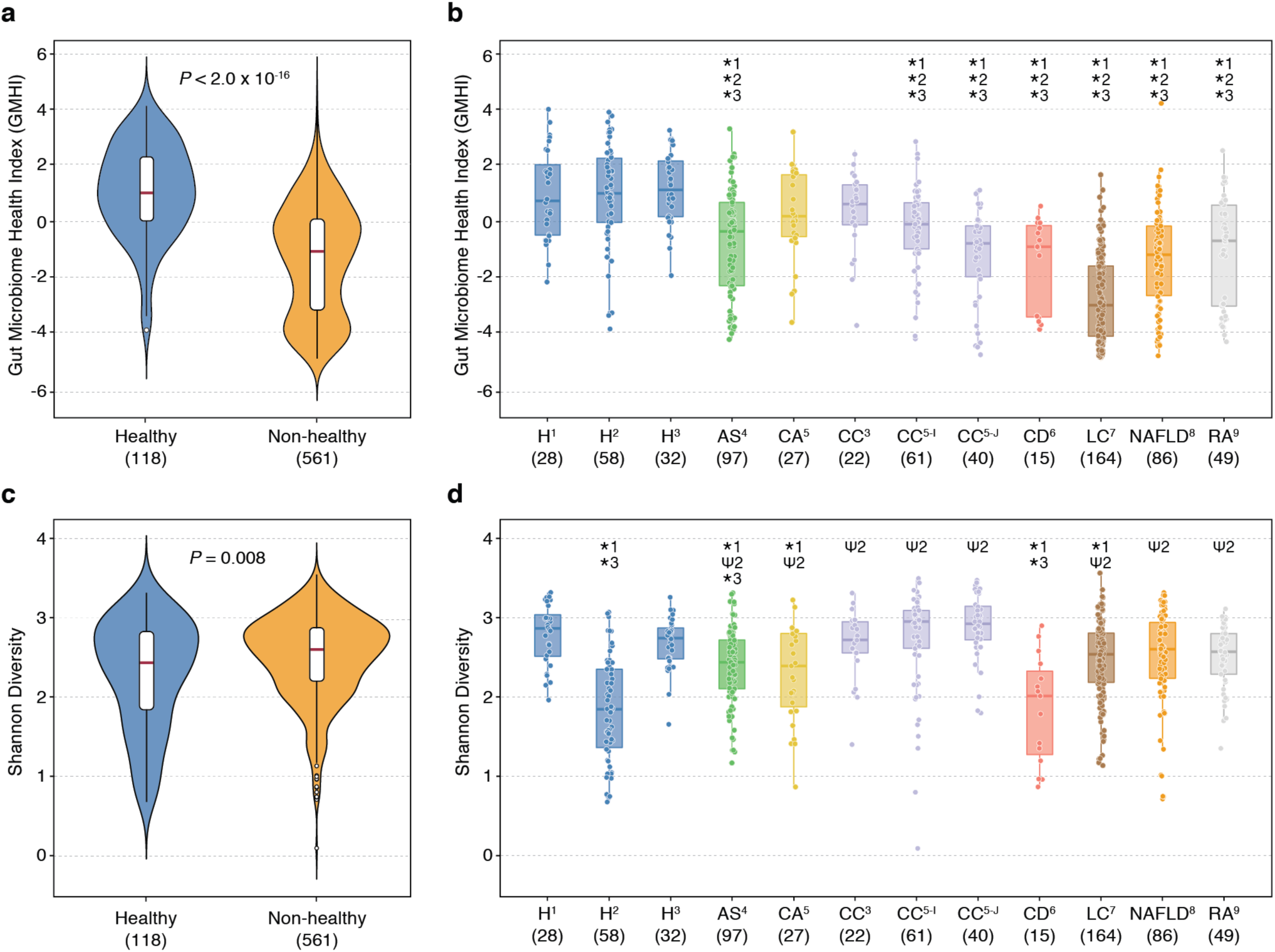
GMHI demonstrates strong reproducibility on validation datasets and outperforms Shannon diversity. To validate performance in stratifying healthy and non-healthy phenotypes, GMHI and Shannon diversity were tested on 679 stool metagenome samples independent of the training dataset. Validation datasets consisted of twelve total cohorts ranging across eight healthy and non-healthy phenotypes from nine different studies. **(a)** GMHIs from stool metagenomes of the healthy group (n = 118) were significantly higher than those of the non-healthy group (n = 561). **(b)** All three healthy cohorts (H^1^, H^2^, and H^3^) were found to have significantly higher distributions of GMHI than seven (of nine) non-healthy cohorts (AS^4^, CC^5-I^, CC^5-J^, CD^6^, LC^7^, NAFLD^8^, and RA^9^). No significant differences were found amongst H^1^, H^2^, and H^3^. The number in superscript adjacent to phenotype abbreviations corresponds to a particular study used in validation (see **Supplementary Table 8** for study information). **(c)** Shannon diversity was significantly lower in healthy than in non-healthy individuals. **(d)** Shannon diversity showed very inconsistent results in distinguishing healthy from non-healthy phenotypes: H^1^ had significantly higher distributions than in four non-healthy cohorts (AS^4^, CA^5^, CD^6^, and LC^7^); H^2^ had significantly lower distributions than in eight non-healthy cohorts (AS^4^, CA^5^, CC^3^, CC^5-I^, CC^5-J^, LC^7^, NAFLD^8^, and RA^9^); and H^3^ had significantly higher distributions than in only two non-healthy cohorts (AS^4^ and CD^6^). Moreover, H^1^ and H^3^ were found to have significantly higher Shannon diversity than H^2^. *P*-values shown above the violin plots were found using Mann-Whitney *U* test. * and Ψ indicates significantly higher distribution in healthy and in a non-healthy phenotype, respectively (*P* < 0.01; Mann-Whitney *U* test). The number adjacent to * and Ψ indicates the healthy cohort (H^1^, H^2^, or H^3^) to which the respective cohort was compared. The sample size of each group is shown in parentheses. AS, ankylosing spondylitis; CA, colorectal adenoma; CC, colorectal cancer; CD, Crohn’s disease, H, healthy; LC liver cirrhosis; NAFLD, non-alcoholic fatty liver disease; RA, rheumatoid arthritis.

To investigate more closely GMHI performances on a cohort-by-cohort basis, we examined the twelve total cohorts ranging across eight healthy and non-healthy phenotypes from eight additional published studies and one newly sequenced cohort. As shown in **Fig. 6b**, all three healthy cohorts were found to have significantly higher distributions of GMHI than seven (of nine) non-healthy phenotype cohorts (*P* < 0.01; Mann-Whitney *U* test). The classification accuracies for these three healthy cohorts were 87.5% (28 of 32), 74.1% (43 of 58) and 71.4% (20 of 28); alternatively, the classification accuracies for the non-healthy phenotype cohorts were the following: 94.5% (155 of 164) for LC; 75.6% (65 of 86) for NAFLD; 73.3% (11 of 15) for CD; 67.3% (33 of 49) for RA; 55.7% (54 of 97) for AS; 37.0% (10 of 27) for CA; and 77.5% (31 of 40), 47.5% (29 of 61), and 27.3% (6 of 22) for three different cohorts of CC. Strikingly, GMHI performed well (>75.0%) in predicting adverse health for LC and NAFLD, although stool metagenomes from patients with liver disease were not part of the original training data. This finding suggests that GMHI could be applied beyond the original twelve phenotypes (of the non-healthy group) used during the index training process. Overall, the remarkable reproducibility of GMHI implies that the highly diverse and complex features of gut microbiome dysbiosis implicated in pathogenesis were reasonably well captured during the dataset integration and original formulation of GMHI.

In contrast to GMHI, species-level Shannon diversity of the gut microbiome was generally a poor predictor of health when applied to the same validation dataset. Opposite to our expectations, the Shannon diversities of the healthy validation group were slightly lower than those of the non-healthy validation cohort (*P* = 8.1×10^−3^, Mann-Whitney *U* test; **Fig. 6c**). Across the individual cohorts, Shannon diversity demonstrated a far less robust and consistent stratification between healthy and non-healthy compared to GMHI (**Fig. 6d**): the first healthy cohort was found to have significantly higher Shannon diversity than four disease cohorts (AS, CA, CD, and LC); the second healthy cohort was found to actually have significantly lower Shannon diversity than eight disease cohorts (AS, CA, three cohorts of CC, LC, NAFLD, and RA); and the third healthy cohort was found to have significantly higher Shannon diversity than only two disease cohorts (AS and CD) (*P* < 0.01, Mann-Whitney *U* test). Additionally, two healthy cohorts were found to have significantly higher distributions of Shannon diversity than the third (*P* < 0.01, Mann-Whitney *U* test), whereas no significant differences were found amongst all three healthy cohorts for GMHI. Furthermore, the highest Shannon diversities were observed in two colorectal cancer cohorts, whereas the highest GMHIs were observed in the three healthy cohorts. Finally, as was the case with Shannon diversity, we found that 80% abundance coverage and species richness also displayed very weak and inconsistent stratification between healthy and non-healthy cohorts (**Supplementary Fig. 6**).

## DISCUSSION

Given the growing body of evidence linking alterations in the gut microbiome to major illnesses, the lack of analytical tests or rigorous quantitative methods that provide clear insight into the health status of one’s gut microbiome is a critical issue. Moreover, microbiome data is highly complex with enormous sample-level heterogeneity. In these regards, providing a simple measure to quantify the degree of wellness, or the divergence away from a healthy condition, would address a clinical need in current microbiome science. Herein, we address this challenge accordingly: i) by integrating massive amounts of publicly available data (4,347 publicly-available, shotgun metagenomic data of gut microbiomes from 34 published studies), we identified a small consortium of 50 microbial species associated with human health (**Table 2**). More specifically, 7 and 43 species were abundant and scarce, respectively, in the healthy group compared to the non-healthy group; ii) we developed the GMHI that determines general health status (i.e., good or adverse health) from a stool specimen. GMHI is a biologically-interpretable, quantitative metric formulated based on the balance between the collective abundances of Health-prevalent and Health-scarce species. In regards to classification performance, GMHI distinguished healthy from non-healthy (as well as individual disease cohorts in intra-study comparisons) far better than methods adopted from ecological principles (e.g., α-diversity indices), thereby paving a path forward to evaluate human (gut microbiome) health through stool metagenomic profiling. Notably, this framework can be applied to other niches of the human body, e.g. quantifying health in skin or oral microbiomes; and iii) on independent validation datasets, we demonstrated the potential of GMHI to distinguish between health and disease, and found strong prediction results for healthy individuals, and cohorts with auto-immune disorders and liver disease. The strong reproducibility in predictive performance on a validation dataset suggests that sufficient dataset integration across a large population could lead to robust predictors of health. This may be due, in part, to the signature encompassing more of the heterogeneity across various sources and conditions, while amplifying signal (against noise) from the repeated phenotype characteristics.

Despite the strong reproducibility in classification performance demonstrated on the validation datasets, more is left to be desired in regards to achieving higher accuracies. We conjecture that misclassifications were partly because of the complex^13,14,48^, stochastic^49,50^, and highly personalized^22,51^ nature of gut microbiome ecologies; all of which complicate the identification and validation of a reliable and robust signature of health. In addition, we cannot rule out the possibility of inaccurate diagnoses for the original phenotype labels. Furthermore, sample collection and processing procedures, laboratory personnel, study run-dates, reagent sources, and measurement instruments are tremendously challenging to control in population-scale investigations. In the long term, in order to find robust gut microbiome-based diagnostic or predictive markers, we envision integrating even larger data repositories to take into consideration more sources of heterogeneity. In turn, as the sample size increases, we expect to obtain a more accurate metric for determining one’s health status based on her/his gut microbiome, as well as new, holistic insights not offered by smaller, individual studies.

Several limitations of our study should be noted when interpreting our results. First, as the stool metagenomes were collected from over 40 published studies, we cannot entirely exclude experimental and technical inter-study batch effects, as is the case for any meta-analysis^30,52^. Our efforts to curtail these batch effects include: i) consensus preprocessing, i.e., downloading all raw shotgun metagenomes (.fastq files) and re-processing each sample uniformly using identical bioinformatics methods; ii) using frequencies of a signal (i.e., prevalence of ‘present’ microbes) as a measure to identify significant associations, rather than comparing or averaging effect-sizes between populations, or performing dataset transformation that may lead to spurious conclusions^53^; and iii) validating the reproducibility of GMHI on independent datasets. Second, given our selection criteria and reasoning (**Methods**), our study does not include all publicly available stool shotgun metagenomic studies and samples. Certainly, more studies and samples can be taken into consideration under more relaxed criteria. Third, in an effort to be as precise and robust as possible in describing taxonomic features of the human gut, our metagenomic analyses were performed using species-level abundances; however, microbial *strains* are clearly the most clinically informative and actionable unit^54,55^. Moreover, different strains within the same species can have significantly different associations with health or disease^56–59^, which could not be considered in our study. Nevertheless, our shotgun metagenomic approach is a significant advancement over 16s rRNA gene amplicon sequencing, which are known to be mostly limited to genus-level investigations^60,61^. Fourth, in the non-healthy group, we pooled samples from only twelve disease or abnormal bodyweight conditions. Certainly, many more pathological states have been linked to the gut microbiome, including neuro-degenerative and psychiatric disorders^62–64^. To truly gain robust insight into a gut microbiome associated with, or predictive of, general wellness, future studies will need to continuously update and expand our findings by encompassing a much broader range of conditions as new data become available. Fifth, we did not answer the question of whether the Health-prevalent microbes were drivers of good health or were a mere reflection; similarly, we did not investigate whether Health-scarce microbes could initiate or mediate disease, or were a common response to adverse health conditions. Although these questions were outside the scope of this current study, future investigations could certainly examine the role of these microbes by carrying out experiments on animal models. By understanding their protective or pathophysiological roles, we speculate that these microbial species could serve as a basis to evaluate the outcome of future clinical interventions. Sixth, we did not consider metagenomic functional profiles to define gut ecosystem health as demonstrated extensively by others^10,65^, as this too was outside the scope of our study. Interestingly, a recent stool metagenomic meta-analysis of functional signatures revealed specific functions that robustly stratify disease individuals from healthy controls^31^. For microbiomes of any phenotype of interest, we posit that analyzing both taxonomic composition and functional potential are both important and complementary directions. Lastly, the longitudinal stability of GMHI remains yet an unanswered question. Testing GMHI within the same individual over time, or assessing the variability with repeat measurements on an individual whose health status is not changing, could be an interesting avenue for future studies.

The discovery of potentially actionable information using computational approaches on high-throughput datasets could bring gut microbiome science one step closer for use in the clinical setting. Indeed, translating microbiome research into health benefits will require a deeper (and more quantitative) understanding of the healthy state. To this end, GMHI could play a significant role in the development of stool metagenomic profiling tests for dynamically monitoring and predicting gut wellness. Furthermore, as the long-term safety and efficacy of fecal microbiota transplantation (FMT) to treat patients remains a concern^66,67^, the widespread success of FMT is highly dependent upon the accurate screening of healthy donors despite their lack of overt disease. In this regard, we anticipate the emerging scenario wherein GMHI could aid the proper selection of healthy donors and stool samples. To this point, GMHI could further facilitate future efforts to define design-principles of a healthy gut microbiome and to establish rational strategies for restoring or maintaining wellness in the form of probiotic ‘cocktail’ communities^68,69^.

## METHODS

### Multi-study integration of human stool metagenomes

We performed exhaustive keyword searches (e.g., “gut microbiome”, “metagenome”, “whole genome shotgun”) in PubMed and Google Scholar for published studies with publicly available whole-genome shotgun (WGS) metagenome data of human stool (gut microbiome) and corresponding subject meta-data (as of March 2018). In studies wherein multiple samples were taken per individual across different time-points, we included only the first or baseline sample in the original study. We excluded studies pertaining to diet or medication interventions, or those with fewer than 10 samples. Samples from subjects who were less than 10 years of age were also excluded from our analysis. Lastly, samples that were collected from disease controls, but were not reported as healthy nor had any mentioning of their health status in the original study, were excluded from our analysis. Raw sequence files (.fastq) were downloaded from the NCBI Sequence Read Archive (SRA) and European Nucleotide Archive (ENA) databases (**Supplementary Table 1**) for the study analysis.

### Re-classification of healthy samples based on reported BMI

Healthy individuals, regardless of whether they had been determined as healthy in the original studies, were considered to be part of the non-healthy group if their reported BMI fell within the range of underweight (BMI < 18.5), overweight (BMI ≥ 25 & < 30), or obese (BMI ≥ 30). Stool metagenome samples from such individuals were re-classified as underweight, overweight, or obese in our analysis.

### Quality control of sequenced reads

Sequence reads were processed with the KneadData quality-control pipeline (http://huttenhower.sph.harvard.edu/kneaddata), which uses Trimmomatic v0.36 and Bowtie2 v0.1 for removal of low-quality read bases and human reads, respectively. Trimmomatic v0.36 was run with parameters SLIDINGWINDOW:4:30, and Phred quality scores were thresholded at ‘<30’. Illumina adapter sequences were removed, and trimmed non-human reads shorter than 60 bp in nucleotide length were discarded. Potential human contamination was filtered by removing reads that aligned to the human genome (reference genome hg19). Furthermore, stool metagenome samples of low read count after quality filtration (< 1M reads) were excluded from our analysis.

### Species-level taxonomic profiling

Taxonomic profiling was done using the MetaPhlAn2 v2.7.0 phylogenetic clade identification pipeline^36^ using default parameters. Briefly, MetaPhlAn2 classifies metagenomic reads to taxonomies based on a database (mpa_v20_m200) of clade-specific marker genes derived from ∼17,000 microbial genomes (corresponding to ∼13,500 bacterial and archaeal, ∼3,500 viral, and ∼110 eukaryotic species).

### Sample-filtering based on taxonomic profiles

After taxonomic profiling, the following stool metagenome samples were discarded from our analysis: i) samples composed of more than 5% unclassified taxonomies (100 samples); and ii) phenotypic outliers according to a dissimilarity measure. More specifically, Bray-Curtis distances were calculated between each sample of a particular phenotype and a hypothetical sample in which the species’ abundances were taken from the medians across those samples. A sample was considered as an outlier, and thereby removed from further analysis, when its dissimilarity exceeded the upper and inner fence (i.e., > 1.5 times outside of the interquartile range above the upper quartile and below the lower quartile) amongst all dissimilarities. This process removed 67 metagenome samples.

### Species-removal based on taxonomic profiles

As taxonomic assignment based on clade-specific marker genes may be problematic for viruses^70,71^, we excluded the 298 of viral origin from our analysis. Species that were labeled as either unclassified or unknown (118 species), or those of low prevalence (i.e., observed in < 1% of the samples included in our meta-dataset; 472 species), were also excluded. Eventually, 313 microbial species across 4,347 stool metagenome samples remained in our study for further analysis (**Supplementary Table 2**).

### Principal Coordinate Analysis based on taxonomic profiles

The R packages ‘ade4’ and ‘vegan’ were used to perform Principal Coordinate Analysis (PCoA) ordination with Bray-Curtis dissimilarity as the distance measure on the stool metagenome samples, which were comprised of arcsin-transformed relative abundances of the aforementioned 313 microbial species identified by MetaPhlAn2.

### Shannon diversity, 80% abundance coverage, and species richness identification

The R package ‘vegan’ was used to calculate Shannon diversity (Shannon index) and species richness based on the species abundance profiles for each sample of our meta-dataset. To identify the 80% abundance coverage for a stool metagenome sample, the smallest number of microbial species that comprise at least 80% of the total relative abundance was identified.

### Construction of gene (KEGG ortholog) copy number profiles for Health-prevalent and Health-scarce species

For every species, a reference strain was selected (**Supplementary Table 5**) and its protein sequences were downloaded from HMP-DACC (https://hmpdacc.org/) or NCBI (www.ncbi.nlm.nih.gov/). Then, the protein sequences were mapped onto the Kyoto Encyclopedia of Genes and Genomes (KEGG) database (KEGG Release 90.0, May 1, 2019) to determine their functional KEGG orthologs. Afterwards, gene copy number profiles were constructed for all reference strains.

### Stool sample collection and processing from patients with rheumatoid arthritis

All stool samples were obtained following written informed consent. The collection of biospecimens was approved by the Mayo Clinic Institutional Review Board (#14-000616). Stool samples from patients with rheumatoid arthritis were stored in their house-hold freezer (−20°C) prior to shipment on dry ice to the Medical Genome Facility Research Core at Mayo Clinic (Rochester, MN). Once received, the samples were stored at −80°C until DNA extraction. DNA extraction from stool samples was conducted as follows: Aliquots were created from parent stool samples using a tissue punch, and the resulting child samples were then mixed with reagents from the Qiagen Power Fecal Kit. This included adding 60 uL of reagent C1 and the contents of a power bead tube (garnet beads and power bead solution). These were then vigorously vortexed to bring the sample punch into solution and centrifuged at 18000G for 15 min. From there, the samples were added into a mixture of magnetic beads using a JANUS liquid handler. The samples were then run through a Chemagic MSM1 according to the manufacturer’s protocol. After DNA extraction, paired-end libraries were prepared using 500ng genomic DNA according to the manufacturer’s instructions for the NEB Next Ultra library prep kit (New England BioLabs). The concentration and size distribution of the completed libraries was determined using an Agilent Bioanalyzer DNA 1000 chip (Santa Clara, CA) and Qubit fluorometry (Invitrogen, Carlsbad, CA). Libraries were sequenced at 23-70 million reads per sample following Illumina’s standard protocol using the Illumina cBot and HiSeq 3000/4000 PE Cluster Kit. The flow cells were sequenced as 150 x 2 paired-end reads on an Illumina HiSeq 4000 using the HiSeq 3000/4000 sequencing kit and HiSeq Control Software HD 3.4.0.38. Base-calling was performed using Illumina’s RTA version 2.7.7.

### Code availability

An R script on how to calculate GMHI for a given stool metagenome sample is available at https://github.com/jaeyunsung/GMHI_2020.

### Data availability of rheumatoid arthritis stool metagenomes

Sequences for the dataset containing rheumatoid arthritis stool metagenomes used for GMHI validation have been deposited at NCBI’s SRA resource under BioProject number PRJNA598446. The deposited sequences includes .fastq files for 49 patients with rheumatoid arthritis. Human reads were identified and removed prior to data upload.

## ACKNOWLEDGMENTS

This work was supported in part by the Mayo Clinic Center for Individualized Medicine (to V.G., M.K., U.B., K.C., N.C., and J.S.), and Mark E. and Mary A. Davis to Mayo Clinic Center for Individualized Medicine (J.S.).

## AUTHOR CONTRIBUTIONS

V.G. and J.S. conceived the problem. V.G., M.K., and J.S. designed all analytical methodologies. V.G. and K.C. performed the computational experiments. All authors analyzed the data. V.G. and J.S. wrote the manuscript, with contributions from other authors. J.M.D. is the principal investigator of the Mayo Clinic Rheumatology Biobank, from which new stool samples were collected from patients with rheumatoid arthritis. All authors reviewed and approved the final manuscript.

## COMPETING INTERESTS

The authors (V.G. and J.S.) disclose an intent to file a patent surrounding the GMHI formula along with the accompanying panel of Health-prevalent and Health-scarce species.

