## Supplementary Methods and Figures for "A Novel Index for Predicting Health Status Using Species-level Gut Microbiome Profiling"

for

### SUPPLEMENTARY METHODS

1. **Identification of microbial species more frequently observed in Healthy group $H$ than in Non-healthy group $N$ (and vice versa)**
   **1.1** Let $p_{H,m_i}$ and $p_{N,m_i}$ be the prevalence of microbial species $i$ ($m_i$), i.e., proportion of samples in a given group where $m_i$ is 'present' (or relative abundance $\geq 1.0 \times 10^{-5}$), in group $H$ and group $N$, respectively.
      **1.1.1** Remark: The relative abundances for all detectable species in a microbiome (metagenome) sample sums to 1.
   **1.2** For $m_i$, we identify prevalence fold-change $f_{m_i}^H$ and prevalence difference $d_{m_i}^H$, defined as $\frac{p_{H,m_i}}{p_{N,m_i}}$ and $p_{H,m_i} - p_{N,m_i}$, respectively.
   **1.3** We introduce $\theta_f$ and $\theta_d$, which are minimum thresholds for $f_{m_i}^H$ and $d_{m_i}^H$, respectively. For all detectable species in a microbiome sample, we identify those that satisfy $f_{m_i}^H \geq \theta_f$ and $d_{m_i}^H \geq \theta_d$.
   **1.4** For all species that satisfy the criteria in **1.3**, we include as an element of 'Health-prevalent' species $M_H$, or the set of species more frequently observed in group $H$ than in group $N$.
   **1.5** To identify 'Health-scarce' species $M_N$, or the set of species more frequently observed in group $N$ than in group $H$, we repeat steps **1.2** through **1.4** with the following considerations:
      **1.5.1** For $m_i$, we identify $f_{m_i}^N$ and $d_{m_i}^N$, defined as $\frac{p_{N,m_i}}{p_{H,m_i}}$ and $p_{N,m_i} - p_{H,m_i}$, respectively.
      **1.5.2** We use the same thresholds $\theta_f$ and $\theta_d$ for identifying $M_N$. In this regard, the species that are eventually chosen to compose $M_H$ and $M_N$ are both dependent on $\theta_f$ and $\theta_d$.
      **1.5.3** Finally, all detectable species that satisfy $f_{m_i}^N \geq \theta_f$ and $d_{m_i}^N \geq \theta_d$ are included in $M_N$.

2. **Identification of $\psi_{M_H}$ ($\psi_{M_N}$), i.e., the 'collective abundance' of species in $M_H$ ($M_N$) in a microbiome sample**
   **2.1** Introduction to $\psi_{M_H}$
      **2.1.1** Let $\psi_{M_H}$ be the 'collective abundance' of $M_H$ species in a microbiome sample. As described below, the calculation of $\psi_{M_H}$ takes into consideration: i) richness, i.e., the numeric count of 'present' taxonomies, of $M_H$ species; and ii) their relative abundances. $\psi_{M_H}$ can be formulated as a *product* of these two traits.
   **2.2** Basic assumptions
      **2.2.1** $\psi_{M_H}$ is positively correlated with $R_{M_H}$, or the richness of $M_H$ species. Thus, $\rho(\psi_{M_H}, R_{M_H}) > 0$.
         **2.2.1.1** Remark: Due to the possible large discrepancy between the cardinality (set size) of $M_H$ and that of $M_N$, we use the *proportion* of 'present' $M_H$ species. As such, $R_{M_H}$ in the above assumption is replaced with $\frac{R_{M_H}}{|M_H|}$. Thus, $\rho\left(\psi_{M_H}, \frac{R_{M_H}}{|M_H|}\right) > 0$.
      **2.2.2** $\psi_{M_H}$ is positively correlated with $\langle M_H \rangle$, or the mean abundance of species in $M_H$. Thus, $\rho(\psi_{M_H}, \langle M_H \rangle) > 0$.
         **2.2.2.1** Remark: As it is common in microbiome data to have discrepancies between species' relative abundances to span several orders of magnitude, we choose to use the geometric mean, rather than the arithmetic mean, to represent the mean relative abundance of $M_H$

species. More specifically, we use the Shannon's diversity index, which is a weighted geometric mean (by definition), and is commonly used in ecological contexts. Thus, for simplicity, we assume that $\langle M_H \rangle \approx \sum_{j \in I_{M_H}} |n_j ln(n_j)|$, where $n_j$ is the relative abundance of species $j$ and $I_{M_H}$ is the index set of $M_H$.

**2.3** Overview

**2.3.1** Given **2.2.1** and **2.2.2**, as well as the non-negativity of $\frac{R_{M_H}}{|M_H|}$ and $\sum_{j \in I_{M_H}} |n_j ln(n_j)|$, let $\psi_{M_H} = c_{M_H} \frac{R_{M_H}}{|M_H|} \sum_{j \in I_{M_H}} |n_j ln(n_j)|$, where $c_{M_H}$ is a constant specific to $M_H$.

**2.3.2** Analogously, let $\psi_{M_N} = c_{M_N} \frac{R_{M_N}}{|M_N|} \sum_{j \in I_{M_N}} |n_j ln(n_j)|$, where $c_{M_N}$ is a constant specific to $M_N$.

### 3. Identification of $h_{i,M_H,M_N}$ , i.e., ratio of $\psi_{M_H}$ to $\psi_{M_N}$ in microbiome sample $i$

**3.1** Formally, the log-ratio of $\psi_{M_H}$ to $\psi_{M_N}$ in sample $i$ can be written as

$$h_{i,M_H,M_N} = log_{10}\left(\frac{c_{M_H}\frac{R_{M_H}}{|M_H|}\sum_{j \in I_{M_H}} |n_j ln(n_j)|}{c_{M_N}\frac{R_{M_N}}{|M_N|}\sum_{j \in I_{M_N}} |n_j ln(n_j)|}\right) = log_{10}\left(\frac{\frac{R_{M_H}}{|M_H|}\sum_{j \in I_{M_H}} |n_j ln(n_j)|}{\frac{R_{M_N}}{|M_N|}\sum_{j \in I_{M_N}} |n_j ln(n_j)|}\right) + log_{10}\left(\frac{c_{M_H}}{c_{M_N}}\right)$$

Although $c_{M_H}$ and $c_{M_N}$ varies based on $M_H$ and $M_N$, we assume $\frac{c_{M_H}}{c_{M_N}} \approx 1$ for simplicity of the analysis. Thus, the above equation can be simplified to

$$h_{i,M_H,M_N} = log_{10}\left(\frac{\frac{R_{M_H}}{|M_H|}\sum_{j \in I_{M_H}} |n_j ln(n_j)|}{\frac{R_{M_N}}{|M_N|}\sum_{j \in I_{M_N}} |n_j ln(n_j)|}\right)$$

**3.2** By definition, $|M_H|$ and $|M_H|$ is the highest richness that can be obtained by $M_H$ and $M_N$ species, respectively, in a particular microbiome sample. However, we cannot rule out the possibility that these maximum values are rarely obtained; if so, then consequently, having a larger set size of $M_H$ (or $M_N$) can generally result in a lower distribution of $\frac{R_{M_H}}{|M_H|}$ (or $\frac{R_{M_N}}{|M_N|}$), potentially leading to biases in $h_{i,M_H,M_N}$ when $|M_H| \gg |M_N|$ or $|M_H| \ll |M_N|$. Therefore, the upper limits we can eventually use in replacement of $|M_H|$ and $|M_N|$ in the formula for $h_{i,M_H,M_N}$ should reflect more of what is observed in actual microbiome data, e.g., samples ranked according to the magnitude between observed $R_{M_H}$ and $R_{M_N}$. In this regard, we use the following procedure to find alternative measures for $|M_H|$ and $|M_N|$:

**3.2.1** We identify $R_{M_H}$ and $R_{M_N}$ for all microbiome samples in groups $H$ and $N$.

**3.2.2** We rank-order (sort) all samples consecutively by two criteria: First, by all values of $R_{M_N}$ in ascending order (from lowest to highest); and then, by all values of $R_{M_H}$ in descending order (from highest to lowest). This sorting strategy prioritizes having the highest possible $R_{M_H}$ (but with the constraint of having $R_{M_N} \approx 0$) for the most top-ranked samples; and having the highest possible $R_{M_N}$ (but with the constraint of having $R_{M_H} \approx 0$) for the most bottom-ranked samples.

**3.2.3** Let $k_H$ be the closest integer to 1% of the number of samples in group $H$. As $H$ is composed of 2,636 samples, we let $k_H$ be 26. Analogously, as $N$ is composed of 1,711 samples, we let $k_N$ be 17.

**3.2.4** We denote $|M_H|'$ as the median $R_{M_H}$ from the top $k_H$ samples, and denote $|M_N|'$ as the median $R_{M_N}$ from the bottom $k_N$ samples.

**3.2.5** We replace $|M_H|$ and $|M_N|$ in the above formula for $h_{i,M_H,M_N}$ with $|M_H|'$ and $|M_N|'$, respectively.

**3.3** In summary, the ratio of $\psi_{M_H}$ to $\psi_{M_N}$ in gut microbiome sample $i$ can be written as

$$h_{i,M_H,M_N} = log_{10}\left(\frac{\frac{R_{M_H}}{|M_H|'}\sum_{j\in I_{M_H}}|n_j ln(n_j)|}{\frac{R_{M_N}}{|M_N|'}\sum_{j\in I_{M_N}}|n_j ln(n_j)|}\right)$$

### 4. Average classification accuracy of $h_{M_H,M_N}$

**4.1** The relative abundances of species in $M_H$ and those in $M_N$ for microbiome sample $i$ can be provided as input features for $\psi_{M_H}$ and $\psi_{M_N}$, respectively, and for the calculation of $h_{i,M_H,M_N}$, which in turn can classify sample $i$ as group $H$ (i.e., $h_{i,M_H,M_N} > 0$), group $N$ (i.e., $h_{i,M_H,M_N} < 0$), or neither (i.e., $h_{i,M_H,M_N} = 0$).

**4.2** We test $h_{M_H,M_N}$ on all samples in groups $H$ and $N$ by finding the average classification accuracy $\chi_{M_H,M_N}$, defined as the average of the proportions of group $H$ and $N$ samples that were correctly classified, or

$$\chi_{M_H,M_N} = \frac{P(h_{i,M_H,M_N} > 0|i \in H) + P(h_{i,M_H,M_N} < 0|i \in N)}{2}$$

where $P(h_{i,M_H,M_N} > 0|i \in H)$ is the proportion of samples in $H$ whose $h_{i,M_H,M_N}$s are positive, and $P(h_{i,M_H,M_N} < 0|i \in N)$ is the proportion of samples in $N$ whose $h_{i,M_H,M_N}$s are negative.

### 5. Determination of optimal sets $M_H^\gamma$ and $M_N^\gamma$

**5.1** To find the final, optimal sets of $M_H^\gamma$ and $M_N^\gamma$, we start by considering a range of thresholds $\theta_f$ and $\theta_d$. Every pair of $\theta_f$ and $\theta_d$ gives different sets of $M_H$ and $M_N$, and in turn, different values of average classification accuracy $\chi_{M_H,M_N}$ (**Supplementary Table 3**).

**5.2** We determine the final, optimal sets of $M_H^\gamma$ and $M_N^\gamma$ (and their corresponding $\theta_f^\gamma$ and $\theta_d^\gamma$) to be those that result in the highest average classification accuracy $\chi_{M_H,M_N}^{max}$.

### SUPPLEMENTARY FIGURES

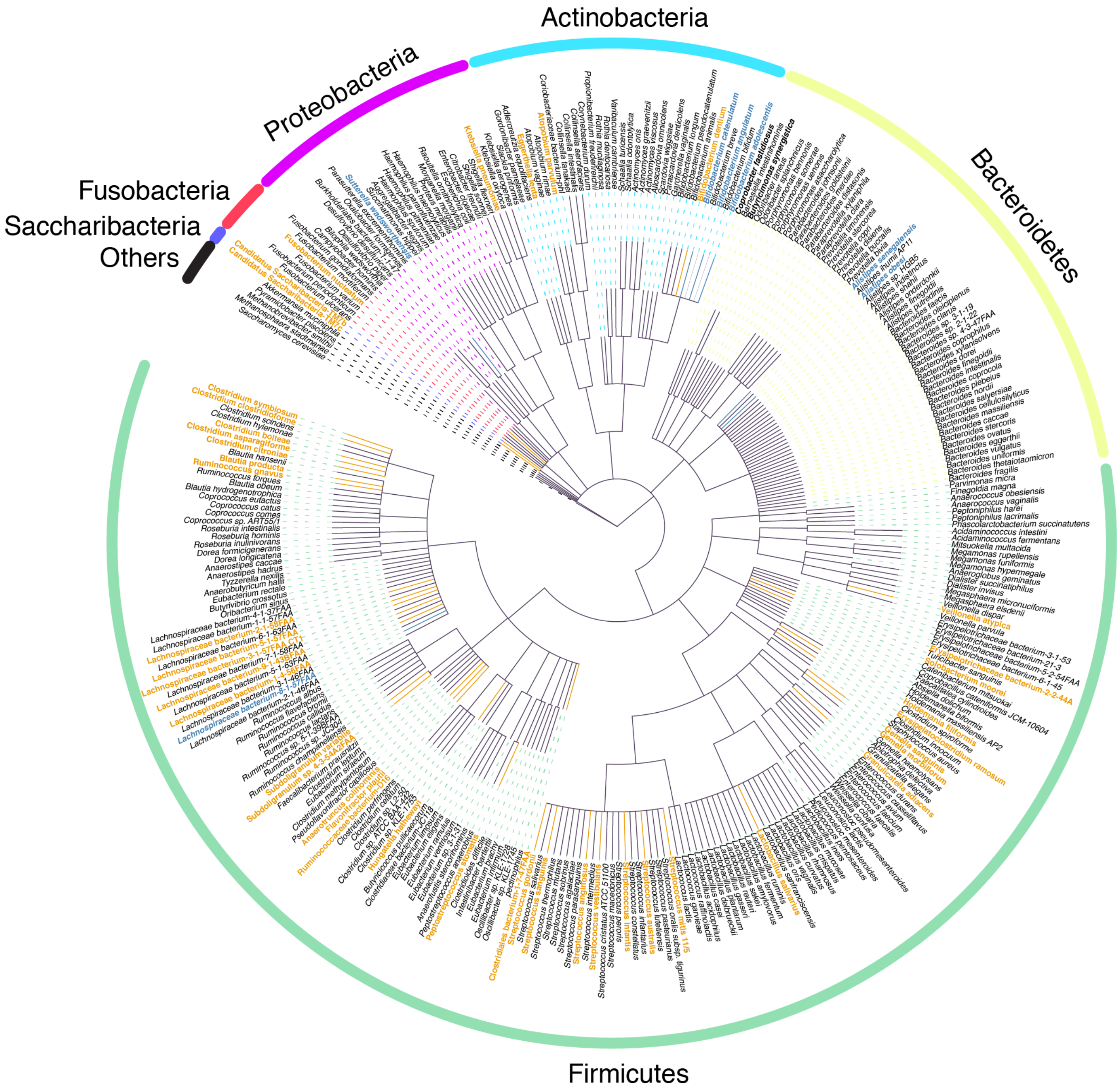


**Supplementary Figure 1. A phylogenetic tree showing the evolutionary relationships among 313 microbial species found to be present across 4,347 stool metagenomes.** Microbial species comprising the Health-prevalent and Health-scarce groups are shown in blue and orange, respectively. Species are grouped according to their phyla (outer circle labels). 'Others' correspond to the Verrucomicrobia (for *Akkermansia muciniphila*), Synergistetes (for *Pyramidobacter piscolens*), Ascomycota (for *Saccharomyces cerevisiae*), and Euryarchaeota (for *Methanobrevibacter smithii* and *Methanosphaera stadtmanae*) phyla.

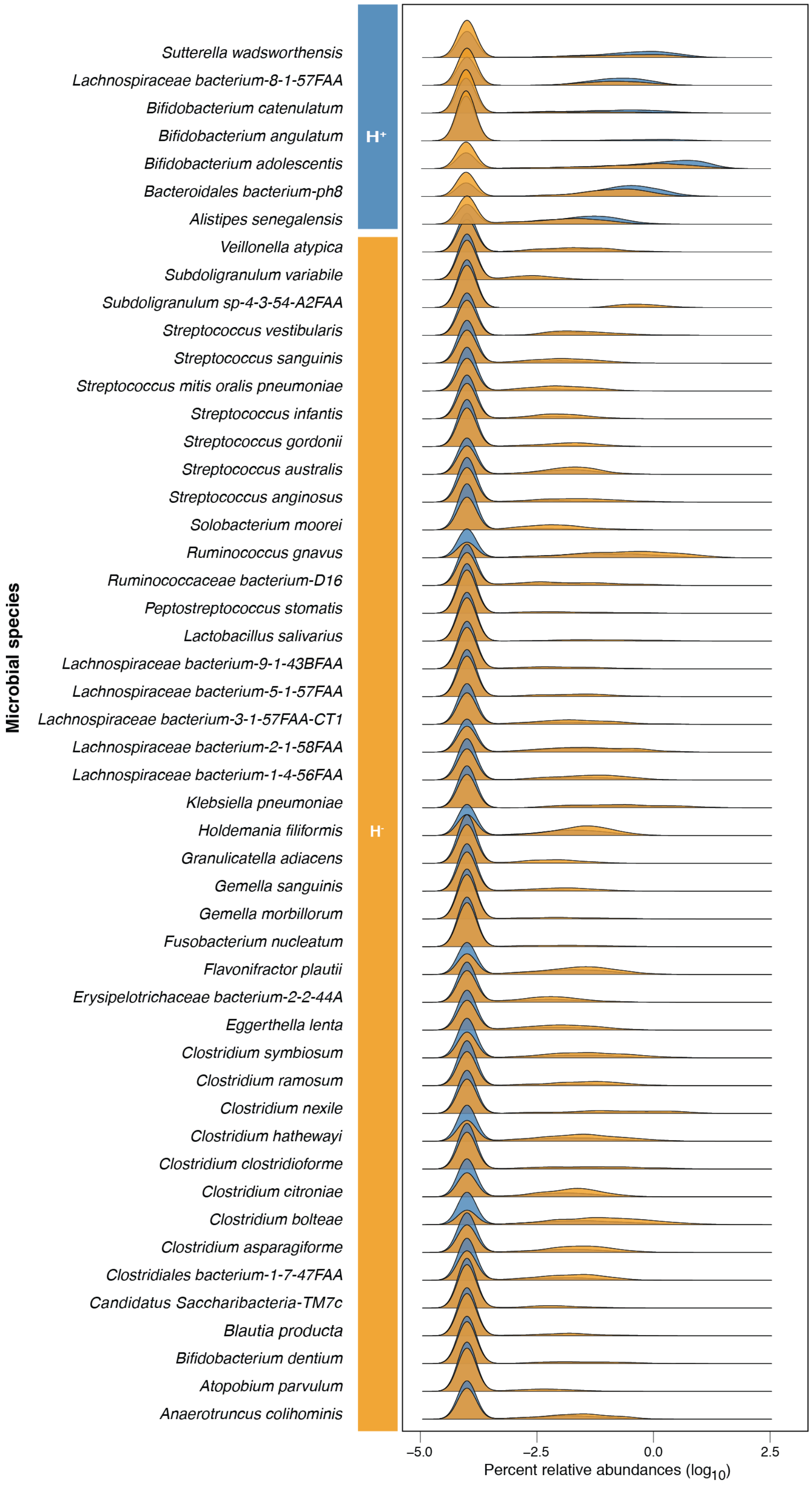


**Supplementary Figure 2. Relative abundance distributions of all 50 species in the healthy (blue density plot) and non-healthy (orange density plot) groups.** Health-prevalent and Health-scarce species show higher relative abundance distributions among healthy and non-healthy gut microbiome samples, respectively.

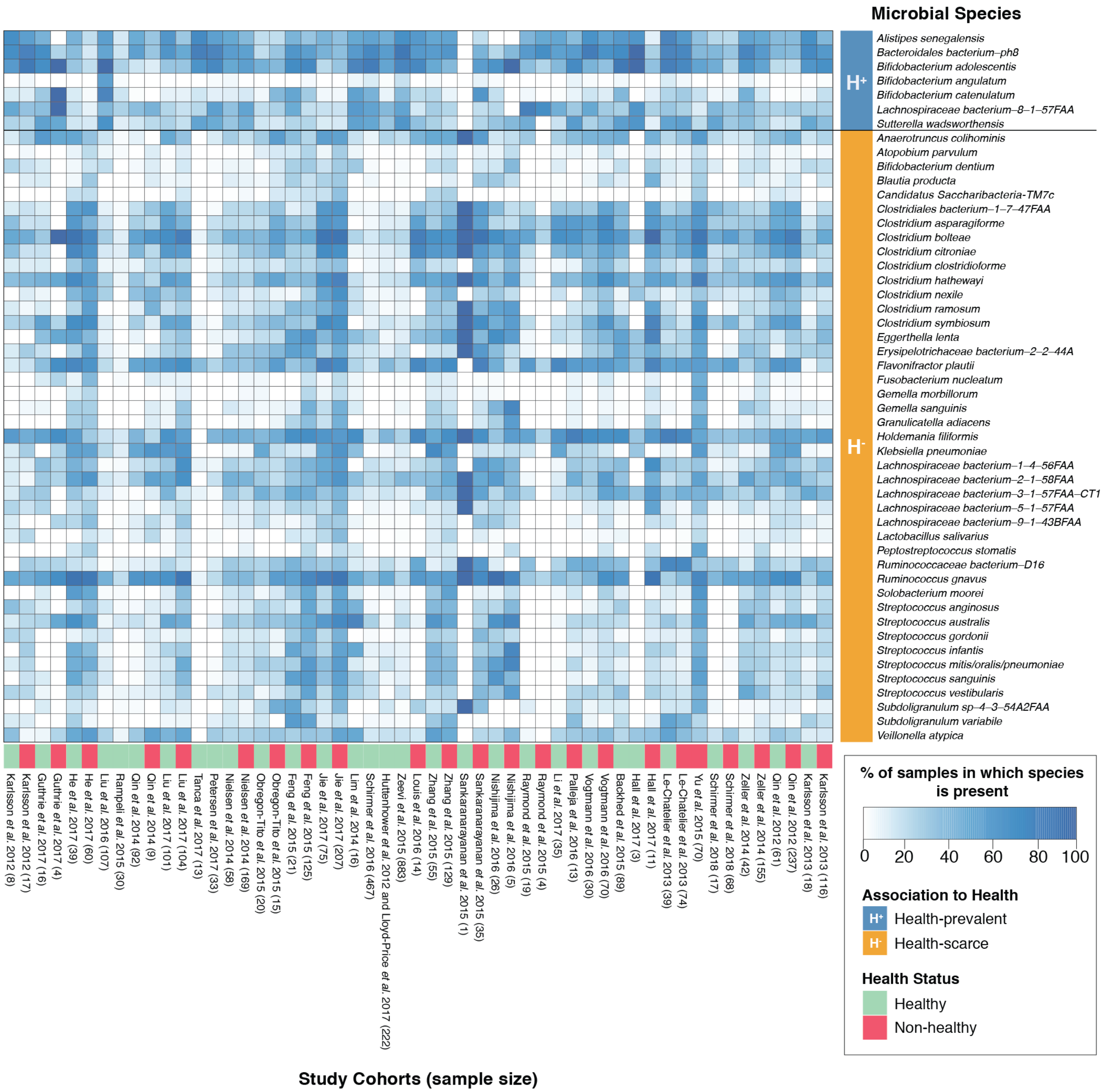


**Supplementary Figure 3.** Heatmap indicating the prevalence of Health-prevalent and Health-scarce species in the healthy and/or non-healthy cohorts from each of the 34 published studies comprising the stool metagenome meta-dataset.

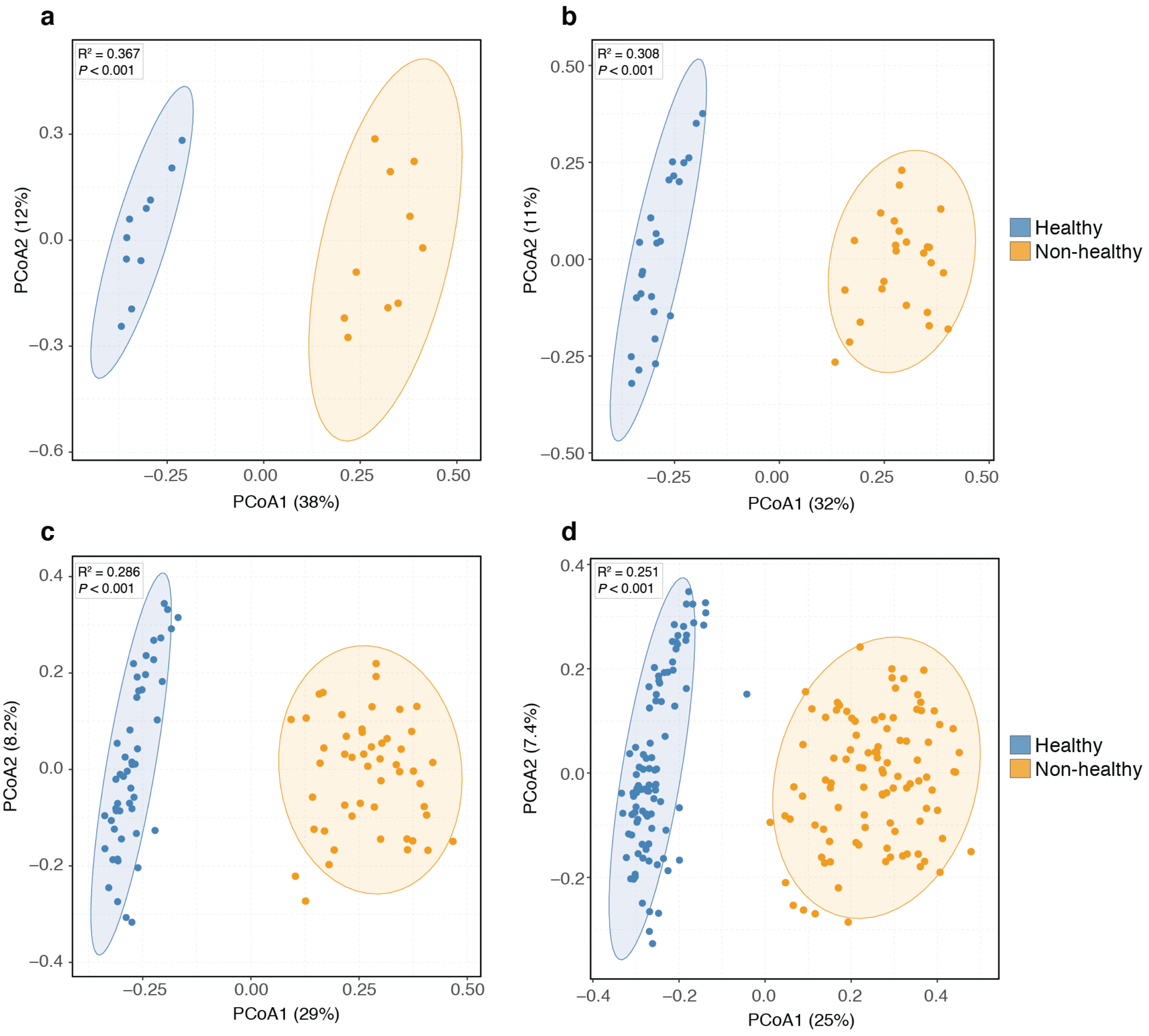


**Supplementary Figure 4.** Top healthy and non-healthy stool metagenomes, as defined by their GMHIs, show clear separation based on gut microbiome composition. Principal Coordinates Analysis (PCoA) ordination plot based on Bray-Curtis distances for top **(a)** 10; **(b)** 25; **(c)** 50; and **(d)** 100 healthy and non-healthy stool metagenomes samples (identified based on GMHI score) show that healthy and non-healthy groups have significantly different distributions of gut microbiome profiles ($P$ < 0.001, PERMANOVA). Each point corresponds to a sample. Ellipses correspond to 95% confidence regions.

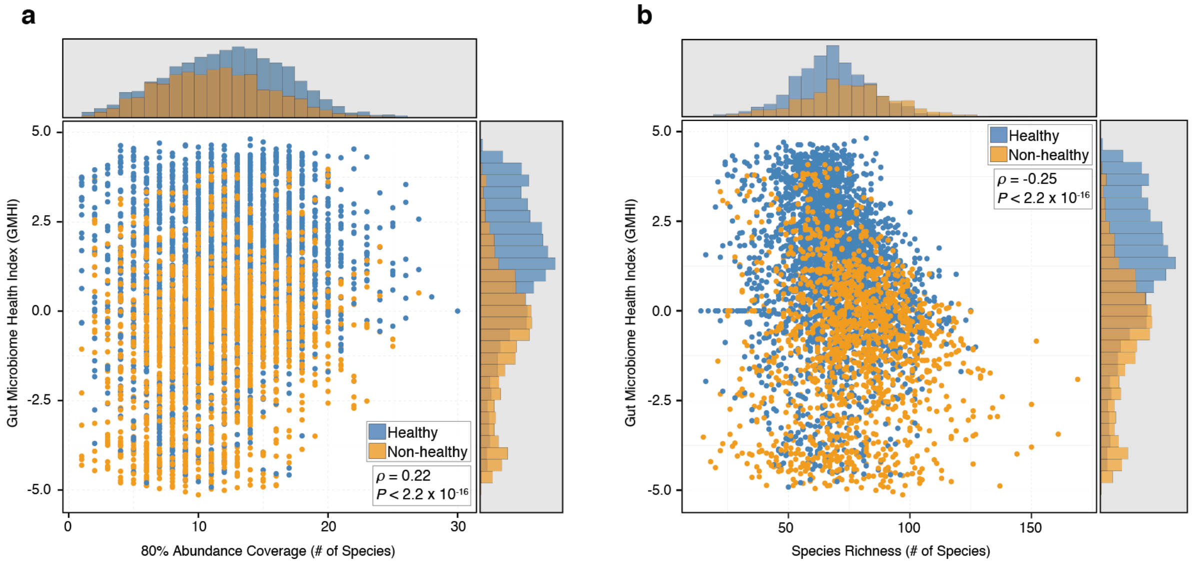


**Supplementary Figure 5.** GMHI stratifies healthy and non-healthy groups more strongly than **(a)** 80% abundance coverage; and **(b)** species richness. Each point in the scatter-plot corresponds to a sample. Histograms show the distribution of healthy (blue) and non-healthy (orange) samples based on the parameter of each axis. In general, GMHI demonstrates weak correlations with 80% abundance coverage (Spearman's $\rho = 0.22$, $P < 2.2\times10^{-16}$) and richness (Spearman's $\rho = -0.25$, $P < 2.2\times10^{-16}$).

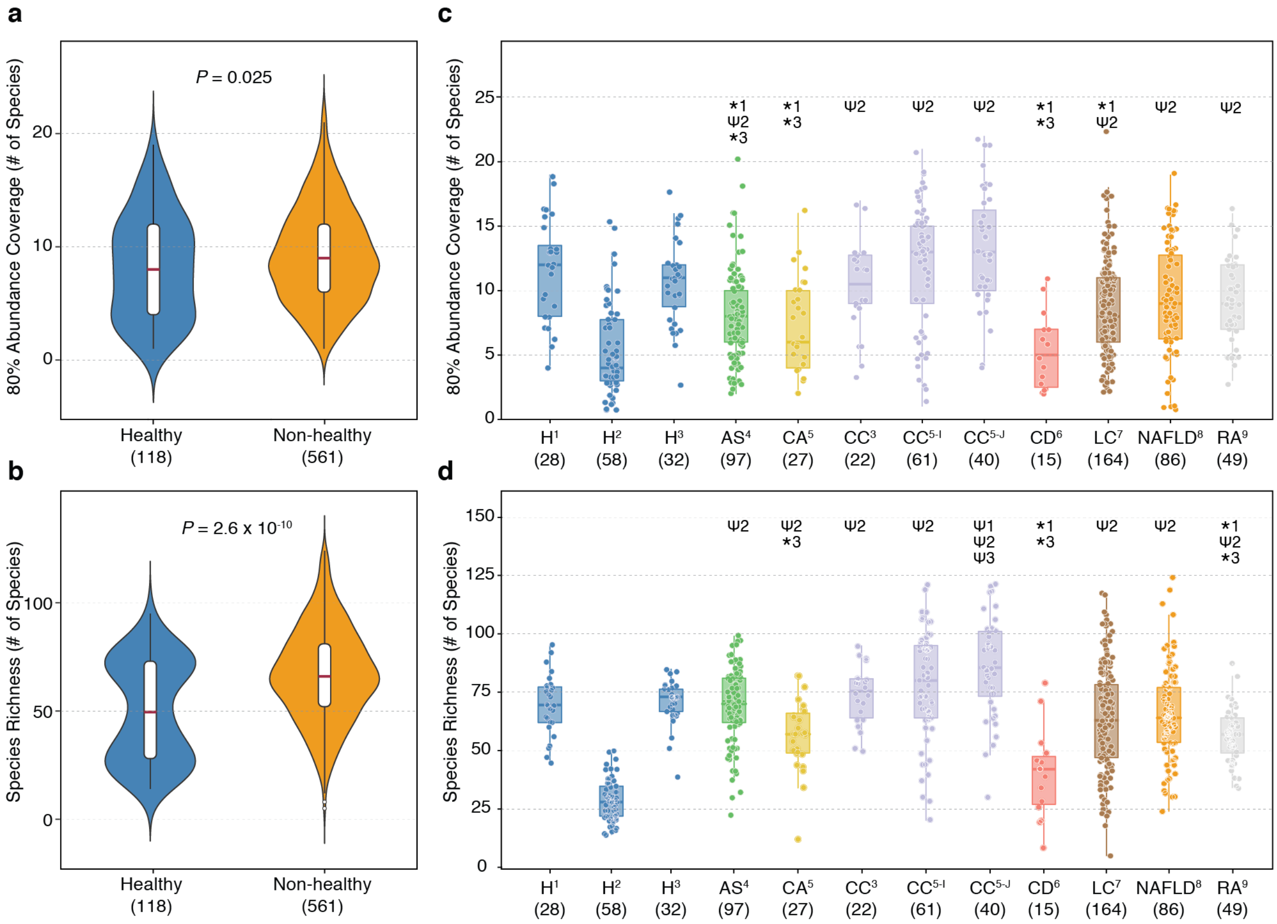


**Supplementary Figure 6. 80% abundance coverage and species richness do not consistently distinguish healthy from non-healthy phenotypes. (a)** 80% abundance coverage and **(b)** species richness in stool metagenomes are significantly different between the healthy group (n = 118) and the non-healthy group (n = 561). **(c)** 80% abundance coverage showed very inconsistent results in distinguishing healthy from non-healthy phenotypes: $H^{1}$ had significantly higher distributions in four non-healthy cohorts ($AS^{4}$, $CA^{5}$, $CD^{6}$, and $LC^{7}$); $H^{2}$ had significantly lower distributions in seven non-healthy cohorts ($AS^{4}$, $CC^{3}$, $CC^{5\text{-}I}$, $CC^{5\text{-}J}$, $LC^{7}$, $NAFLD^{8}$, and $RA^{9}$); and $H^{3}$ had significantly higher distributions in only three non-healthy cohorts ($AS^{4}$, $CA^{5}$, and $CD^{6}$). **(d)** Similarly, species richness also showed inconsistency in distinguishing healthy from non-healthy phenotypes: $H^{1}$ had significantly higher distributions in two non-healthy cohorts ($CD^{6}$, and $NAFLD^{8}$) and lower distribution in one non-healthy cohort ($CC^{5\text{-}J}$); $H^{2}$ had significantly lower distributions in seven non-healthy cohorts ($AS^{4}$, $CC^{3}$, $CC^{5\text{-}I}$, $CC^{5\text{-}J}$, $LC^{7}$, $NAFLD^{8}$, and $RA^{9}$); and $H^{3}$ had significantly higher distributions in three non-healthy cohorts ($CA^{5}$, $CD^{6}$, and $RA^{9}$) but lower distribution in one non-healthy cohort ($CC^{5\text{-}J}$). *P*-values shown above the violin plots were found using Mann-Whitney *U* test. * and Ψ indicates significantly higher distribution in healthy and in a non-healthy phenotype, respectively ($P < 0.01$; Mann-Whitney *U* test). The number adjacent to * and Ψ indicates the healthy cohort ($H^{1}$, $H^{2}$, or $H^{3}$) to which the respective cohort was compared. The sample size of each group is shown in parentheses. AS, ankylosing spondylitis; CA, colorectal adenoma; CC, colorectal cancer; CD, Crohn's disease, H, healthy; LC liver cirrhosis; NAFLD, non-alcoholic fatty liver disease; RA, rheumatoid arthritis.
